# Differential Enhancer Activity and FOXF1 Levels Contribute to Higher Inflammatory Gene Expression of Fetal/Neonatal Versus Adult Fibroblasts in IR-induced Senescence

**DOI:** 10.64898/2026.06.24.734246

**Authors:** Reem Hamed, Régis Courbeyrette, Alexander G. Foote, Susan L. Thibeault, Nicolas O. Fortunel, Laure Crabbe, Carl Mann

## Abstract

Some key inflammatory genes controlled by the RELA transcription factor are thought to be highly expressed in fibroblasts induced into senescence by ionizing radiation (IR) as part of the Senescent-Associated Secretory Phenotype (SASP). However, this view is based largely on studies of a limited number of fibroblast cell lines derived from fetal lung or neonatal foreskin. Here, we show that more than half of the primary adult fibroblast strains examined exhibit only weak induction of RELA-dependent inflammatory genes following IR-induced senescence. We define these fibroblasts as “low-responding” to distinguish them from fibroblasts that express high levels of inflammatory gene expression in response to IR. RNA-seq analysis indicated particularly weak IL1A and IL1B expression in low-responding fibroblasts. IL1-alpha and IL1-beta participate in a positive amplification loop for inflammatory gene expression in senescence. Addition of recombinant IL1-alpha or IL1-beta to these fibroblasts sufficed to induce high expression of inflammatory genes. Low-responding fibroblasts thus exhibit cell-autonomous defects in IL1A and IL1B gene activation in response to IR that explains their overall low expression of RELA-targeted inflammatory genes. This defect was correlated with reduced chromatin accessibility and H3-K27-acetylation at 2 putative enhancers in the intergenic region separating IL1A and IL1B, and deletion of either of these enhancers inhibited inflammatory gene expression in IR-induced senescence. Fibroblasts express distinct transcriptomes and we found that differential expression of the FOXF1 transcription factor gene in high-responding WI38 fetal lung fibroblasts contributes to inflammatory gene expression after IR. Our observations indicate that fibroblasts can be distinguished by their ability to manifest cell-autonomous induction of inflammatory genes under conditions of IR-induced senescence.

## Introduction

Cellular senescence is a stress response of mammalian cells associated with a stable arrest of cell proliferation and modifications of molecular profiles, notably at the transcriptome level [1]. This pathway has beneficial effects including wound healing [2] and in tumor suppression, by blocking the proliferation of cells with persistent DNA damage or that express hyper-mitogenic oncogenes [3], but it has also been implicated in aging-related diseases [4]. Many of the adverse effects of cellular senescence have been related to the secretion of inflammatory factors that are part of the senescence-associated secretory phenotype [5]. The activation of inflammatory gene expression in senescence requires several transcription factors including RELA, CEBPB, and the AP1 family of transcription factors [5]. The AP1 family are pioneer transcription factors [6] that open chromatin regions and thereby allow the binding of non-pioneer transcription factors such as RELA that would otherwise be unable to access their binding sites in closed chromatin [7]. RELA is a master activator of cytokine and chemokine genes in both immune and non-immune cells [7]. The SASP was initially proposed to be a highly-conserved phenotype of senescent cells [8], but subsequent studies have shown cell type variability in its composition [5]. Nevertheless, it is still generally accepted that fibroblasts express high levels of inflammatory genes during senescence induced by persistent DNA damage or by the expression of hyper-mitogenic oncogenes [5]. However, this conclusion was inferred from the study of a relatively small number of fibroblast cell lines derived from fetal lung or neonatal foreskin [9]. Of note, our survey of the literature indicated only a small number of studies characterizing adult fibroblasts. The original paper suggesting a wide conservation of the SASP in senescence included a sample of adult breast fibroblasts that expressed lower levels of inflammatory factors compared to fetal/neonatal fibroblasts in response to IR [8]. A second paper also indicated that adult breast fibroblasts did not induce inflammatory genes in response to irradiation [10]. Finally, we found a study in which a strain of normal adult skin fibroblasts did induce inflammatory genes in response to IR [11].

In this study, we show that the majority of primary fibroblasts derived from diverse adult tissues express inflammatory factors at lower levels than fetal/neonatal fibroblasts during senescence induced by IR. We further identify candidate cis-acting enhancers and trans-acting transcription factors that distinguish the inflammatory response of fetal WI38 fibroblasts from adult fibroblasts.

## Results

### Mammary M168 fibroblasts do not express high levels of CXCL8 or IL6 in IR-induced senescence

During phenotypic characterization of a primary mammary fibroblast cell line (M168 cells) induced into senescence by IR, we were surprised to see little or no induction of IL6 and CXCL8 expression in conditions in which similarly-treated fetal lung WI38 fibroblasts showed higher expression of these inflammatory factors as expected (**Fig 1**). In these IR conditions, proliferation (EdU incorporation) and SA-ß-galactosidase assays showed that the majority of the M168 and WI38 cells were induced into senescence (**Fig S1**). M168 cells also showed low inflammatory gene expression in senescence induced by etoposide that creates DNA damage by blocking DNA topoisomerase II on chromatin (**Fig S2**). The majority of both WI38 and M168 cells induced into senescence by DNA damage contained gH2AX foci associated with persistent DNA damage signaling (**Fig S3**). M168 cells that were passaged until replicative senescence that results from DNA damage at eroded telomere sequences also expressed low levels of inflammatory genes (**Fig. S4**). In particular, RT-qPCR experiments showed no significant induction in replicative senescence versus young proliferating cells for the IL1A, IL1B, and CXCL1/2/3/5/6/8 genes, although the IL6 gene was significantly induced (**Fig. S4E**). Thus, a distinct pathway for IL6 induction independent of other RELA-target genes must exist during replicative senescence of M168 cells.

**Fig 1.**
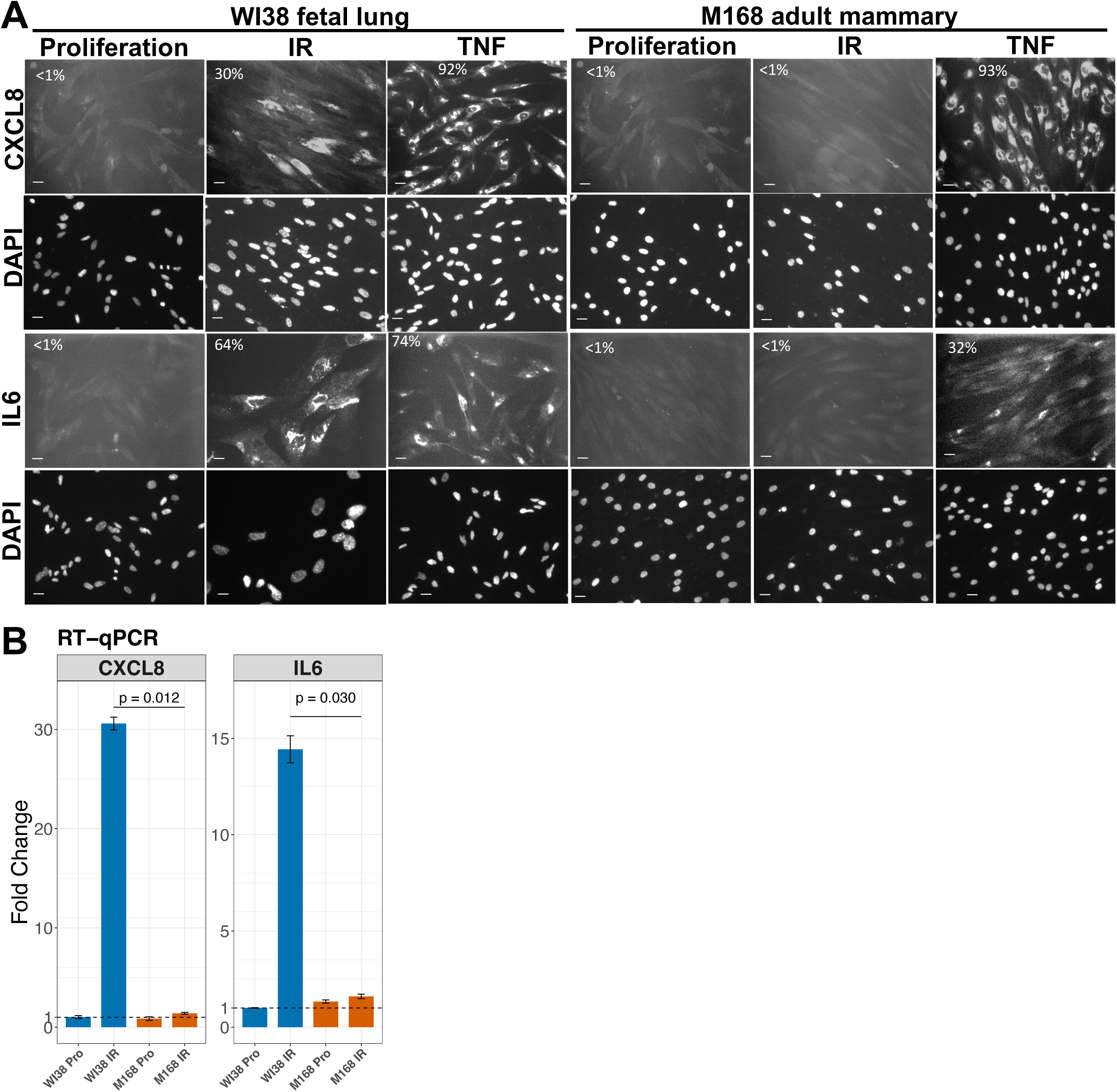
M168 primary mammary fibroblasts express low levels of CXCL8 and IL6 in IR-induced senescence. (**A**) Immunofluorescence analysis of CXCL8 and IL6 expression in WI38 and M168 fibroblasts in control proliferating cells or 10 days after 40 Gy X-irradiation (IR), or after treatment of proliferating cells with 10 ng/ml TNF-alpha for 2 hours (TNF). The percentage of positive cells is indicated after scoring a minimum of 100 cells. The white scale bar is 10 µm. (**B**) RT-qPCR analysis of CXCL8 and IL6 RNA levels under the indicated conditions (mean +/- SEM, n=3, Welch’s t-test).

The absence of high RELA-targeted inflammatory gene expression in the M168 cells induced into senescence by DNA damage appears to be an intrinsic feature of the cells that could not be explained by specific in vitro culture conditions. Treatment of the mammary fibroblasts with TNF-alpha led to a rapid and robust inflammatory gene expression (**Fig 1A**), as did expression of the B-RAF-V600E oncogene (**Fig S5**) that promotes a form of oncogene-induced senescence with minimal DNA damage [12]. Immunofluorescence experiments also showed a strong nuclear translocation of RELA and expression of CXCL8 in WI38 and M168 fibroblasts in response to exogenous addition of IL1-beta (**Fig S6**). In contrast, nuclear localization of RELA and CXCL8 expression was slightly weaker in WI38 induced into senescence by IR and much weaker in M168 cells induced into senescence by IR (**Fig S6**). Thus, the weak expression of inflammatory genes in M168 cells in response to IR is correlated with a weak nuclear translocation of RELA.

These results indicated that the mammary fibroblasts studied here were not generally refractory to the activation of inflammatory gene expression, but they were weakly responsive to activation induced by DNA damage. Interestingly, although the initial article describing the SASP suggested that high inflammatory gene expression was a highly conserved feature of senescent cells, perusal of the data shown in the first figure of their article indicated that primary breast fibroblasts in IR-senescence secreted lower levels of inflammatory factors than did fetal/neonatal WI38, IMR90, HCA2, and BJ fibroblasts [8]. We thus decided to test if high inflammatory gene expression in IR-SEN is commonly associated with primary adult fibroblasts from several tissues.

### Most adult fibroblasts express low levels of RELA-activated inflammatory genes in IR-induced senescence

We examined 16 primary adult fibroblast strains (**Table S1**) derived from 11 individuals of diverse ages and sex by RNA-seq analysis to determine whether high levels of inflammatory gene expression are commonly induced during IR-SEN (**Table S2**). Fibroblasts were obtained from the abdomen, breast, lung, gingiva, scalp, trachea, and vocal folds. Cells were X-irradiated with a dose such that at least 80% of cells were induced into senescence as determined by proliferation and senescence-associated ß-galactosidase assays. RNA-seq analysis was performed 10 days post-irradiation. In addition to our WI38 and adult primary fibroblast RNA-seq datasets, we included in our analysis previously published RNA-seq data for HCA2 foreskin fibroblasts [13] and some breast Cancer-Associated Fibroblasts (CAFs) [14]. HCA2 foreskin fibroblasts express high inflammatory genes in IR-induced senescence as do fetal lung fibroblasts [13]. Breast CAFs are of interest because they can constitutively express high levels of inflammatory genes without being senescent [10,14–16]. Principal component analyses (PCA) indicated that the largest amount of variation (PC1) was associated with the tissue origin of the fibroblasts (**Fig 2**). As was described previously [17–19], heat maps of HOX transcription factor gene expression result in almost perfect clustering of the fibroblasts by their tissue origin (**Fig S7**). Furthermore, non-HOX Homeobox gene heat maps also allow substantial clustering by tissue origin (**Fig S8**), although less well than for HOX gene expression. Interestingly, gingival and palatial fibroblasts express low levels of all HOX genes, but instead express a high level of some characteristic non-HOX Homeobox transcription factors. Finally, the collection of the most highly variable transcription factor genes, excluding all Homeobox transcription factors, also allows a good clustering of fibroblasts according to their tissue origin (**Fig S9**). The expression of some of these transcription factor genes is modified by IR, so they represent candidate regulators contributing to potential differences of fibroblasts in their response to irradiation. Overall, this analysis supports the proposal that differences in transcription factor gene expression contribute to the tissue identity of fibroblasts, presumably derived from developmentally-driven epigenetic modifications of regulatory elements controlling transcription factor expression [20].

**Fig 2.**
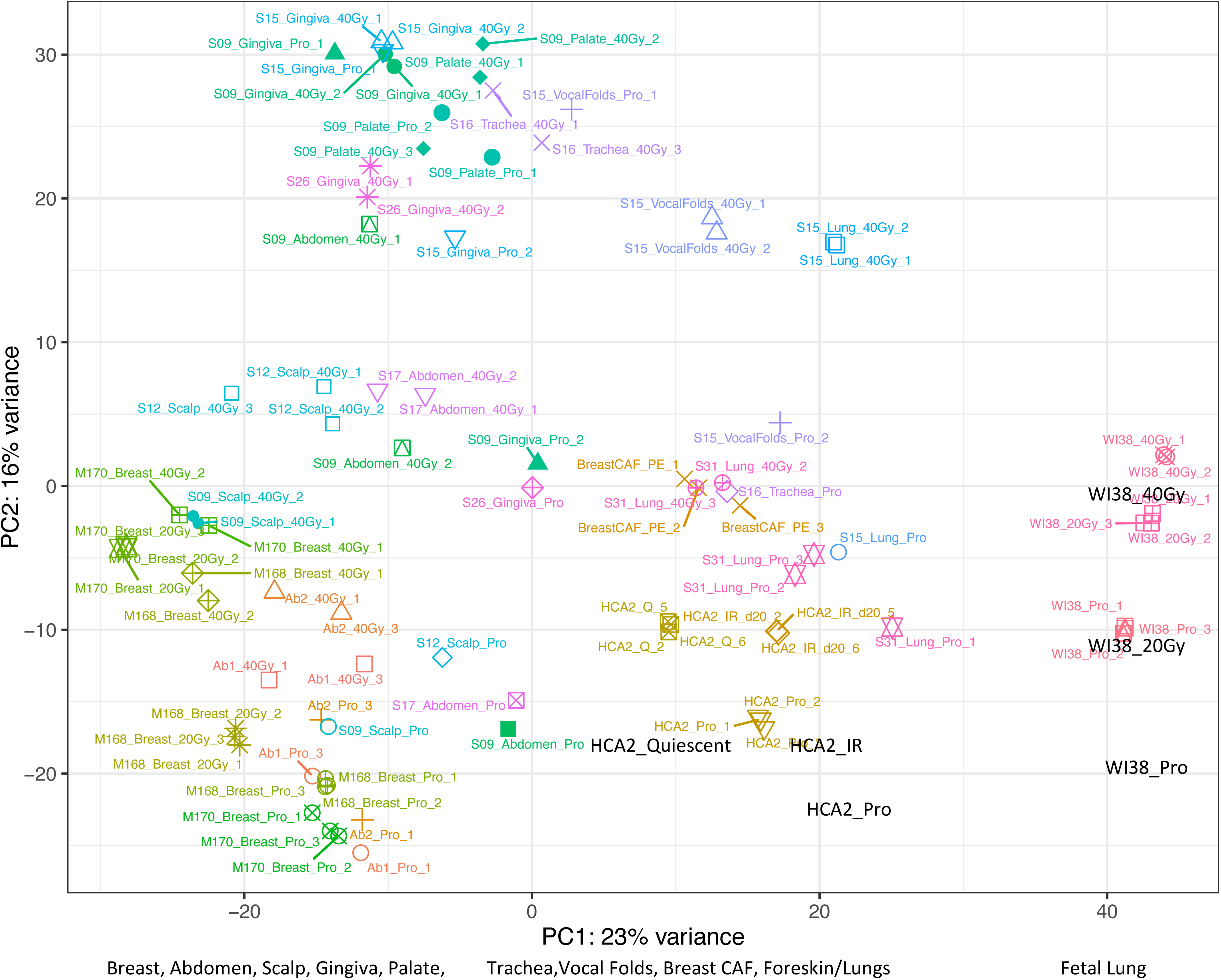
PCA of RNA-seq data for 16 primary adult fibroblasts, WI38 fetal lung and HCA2 neonatal foreskin fibroblasts, and 3 breast cancer fibroblasts. PC1 variation is associated with the tissue origin of the cells and PC2 variation is related to irradiation. The relative position along the PC1 axis of the tissue origin of the fibroblasts is shown below the graph. The HCA2 foreskin fibroblasts included RNA-seq data for samples induced into quiescence by serum starvation (Q) as well as irradiated (IR) and proliferating cells (Pro).

The PC2 axis of variation appears to reflect at least in part whether the fibroblasts were irradiated or not. Irradiated fibroblasts were shifted towards positive loadings relative to the same fibroblasts in proliferation (**Fig 2**). Interestingly, the fetal WI38 and the foreskin HCA2 fibroblasts were mainly segregated from the adult primary fibroblasts by this PCA. WI38 cells irradiated with 40 Gy were our positive control for fibroblasts that express high RELA-activated inflammatory genes in IR-induced senescence. Gene set enrichment analysis (GSEA) of WI38 cells indicated that cell proliferation gene sets were down-regulated by IR as expected for senescent cells, and the only up-regulated gene sets involved inflammatory genes, both RELA-target genes and Interferon-Stimulated Genes (ISGs), as previously reported [21] (**Fig 3A**). We selected 14 RELA-target genes that are highly expressed in WI38 cells in IR-induced senescence to construct a heat map of their expression in all fibroblast lines (**Fig 4A**). The heat map shows that irradiated fetal WI38, irradiated neonatal HCA2, and Breast CAF cells form a cluster in which these genes are highly expressed, whereas all irradiated adult fibroblasts and all proliferating fibroblasts expressed these inflammatory genes at lower levels. Two adult lung fibroblasts and an abdominal fibroblast cell line expressed intermediate (Mid) levels of the RELA-target genes. We also made a heat map of 14 ISGs that are highly expressed in WI38 cells in IR-induced senescence. In this case, about 50% of adult fibroblasts in IR-SEN partially induced ISGs in IR-SEN (Mid), whereas the remaining adult fibroblasts expressed low levels of ISGs and clustered with fibroblasts in proliferation (**Fig 4B**). Thus, RNA-seq analysis revealed that adult fibroblasts in IR-SEN generally express RELA-activated inflammatory genes at lower levels than fetal or neonatal fibroblasts in IR-SEN, but about half of these adult fibroblasts induced ISGs in IR-SEN, although at lower levels compared to fetal/neonatal fibroblasts. GSEA of adult mammary fibroblasts showed down-regulation of cell proliferation gene sets after IR as expected for senescent cells (**Fig 3**). With regards to gene sets that are up-regulated after IR, 2 patterns were observed; fibroblasts such as the mammary M168 showed up-regulation of ISG gene sets, but not NF-kB gene sets, in agreement with the heat map analysis (**Fig 3B**). In contrast, fibroblasts such as the mammary M170 fibroblasts showed no significant up-regulation of ISG or NF-kB gene sets (**Fig 3C**). Indeed, no Hallmark gene sets were significantly up-regulated in these adult fibroblasts in IR-induced senescence.

**Fig 3.**
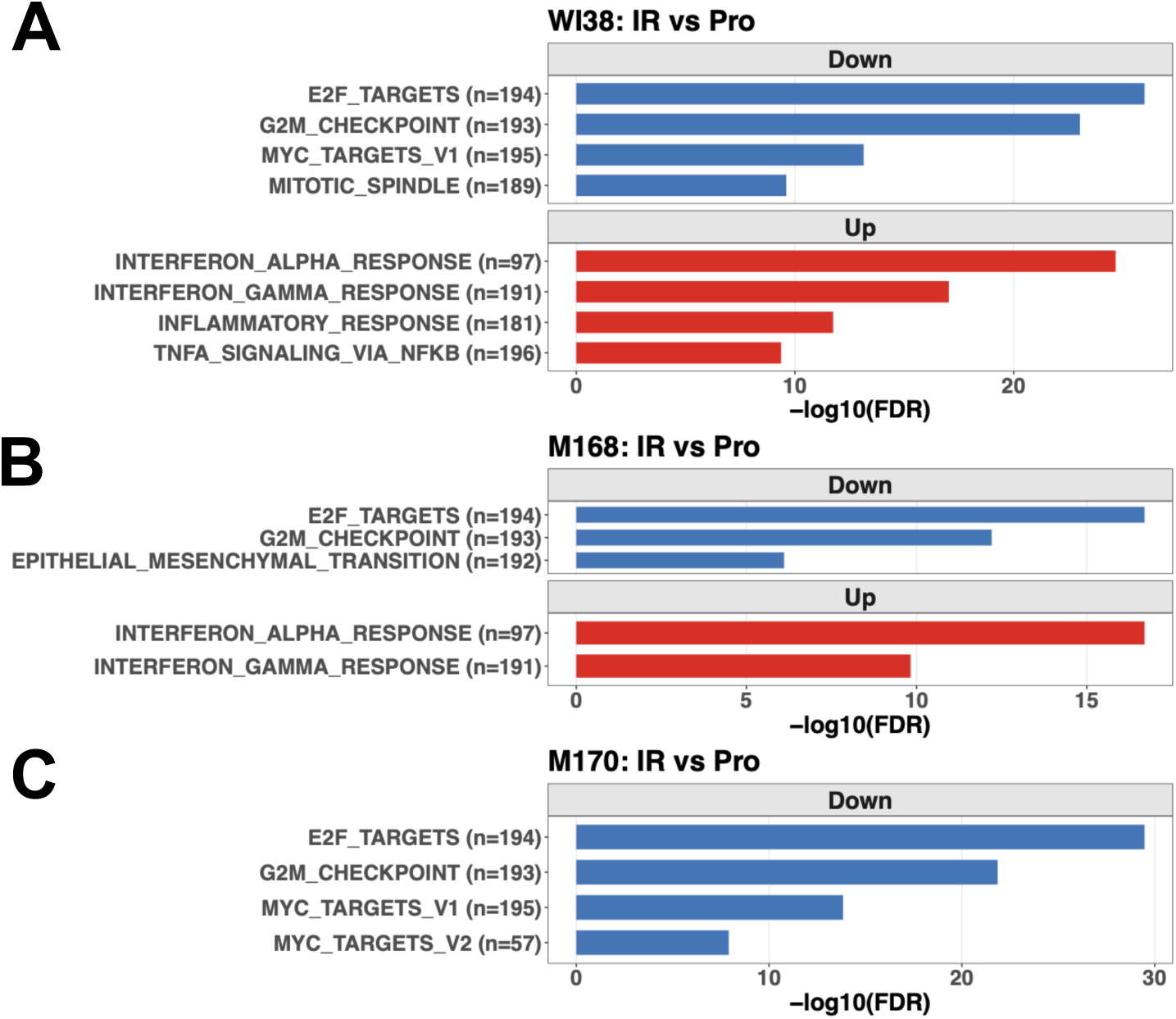
Hallmark gene set enrichment analysis of differentially-expressed genes in cells 10 days after 40 Gy X-irradiation (IR) versus control proliferating cells (Pro). (**A**) WI38 fetal lung fibroblasts. (**B**) M168 adult mammary fibroblasts. (**C**) M170 adult mammary fibroblasts. Shown are all Hallmark gene sets with FDR < 10e-6.

**Fig 4.**
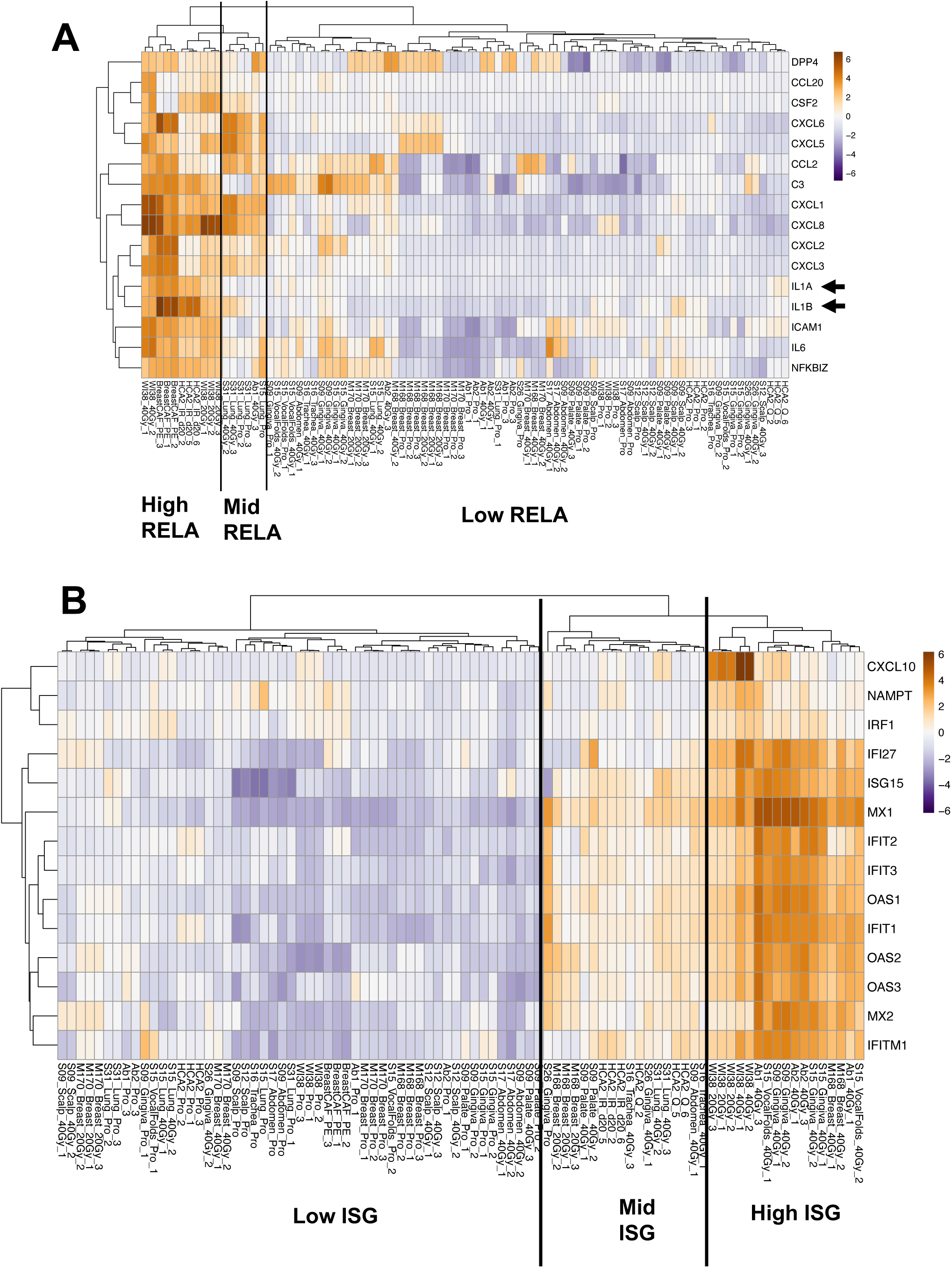
Heat maps of mean-centered gene expression in proliferating and irradiated fibroblasts. (**A**) RELA-target genes that are highly induced in irradiated WI38 cells separated into high, mid, and low expression sample clusters. (**B**) Interferon-stimulatory genes (ISG) that are highly induced in irradiated WI38 cells separated into high mid, and low expression sample clusters. The HCA2 foreskin fibroblasts included RNA-seq data for samples induced into quiescence by serum starvation (Q) as well as irradiated (IR) and proliferating cells (Pro).

### IL1A and IL1B are weakly expressed in adult fibroblasts in IR-SEN and exogenous addition of IL1-alpha or IL1-beta highly activates inflammatory gene expression

We noticed that the IL1A and IL1B genes are weakly expressed in the adult fibroblasts that we examined during proliferation and after IR-induced senescence (**Fig 4A**). This observation drew our attention because IL1-alpha and IL1-beta participate in a positive amplification loop for the activation of RELA in senescent fibroblasts [22–24]. RT-qPCR experiments confirmed that IL1A and IL1B are expressed at lower levels in adult abdominal and mammary fibroblasts compared to fetal WI38 lung fibroblasts in proliferation (**Fig S10A**) or after IR-induced senescence (**Fig S10B,C**). We then tested the effect of adding recombinant IL1-alpha or IL1-beta to fibroblasts in proliferation (**Fig S11**) or 9 days after IR (**Fig S10B,C**). Addition of low levels (4 or 20 pg/ml) of either IL1-alpha or IL1-beta increased the expression of IL1A, IL1B, IL6, and CXCL8 genes in the adult fibroblasts to levels that were similar to that of WI38 fibroblasts (**Fig S10B,C, S11**). This result suggested that a weak induction of IL1A and IL1B in our adult fibroblasts in response to IR may explain the overall weak induction of RELA target genes in these fibroblasts in response to IR. Based on these preliminary dose-response tests by RT-qPCR, we evaluated the global transcriptomic effect of adding 20 pg/ml IL1-alpha for 24 hours to M168 mammary fibroblasts in proliferation and 9 days after IR compared to WI38 cells in proliferation or IR-SEN (**Fig 5 and Table S3**). PCA indicated that PC1 representing the greatest gene expression variation involved fibroblast cell type (WI38 versus M168), PC2 separated proliferating from IR or IL1-alpha treated fibroblasts, and PC3 separated M168_IR from M168_IR+IL1-alpha **(Fig 5A**).

**Fig 5.**
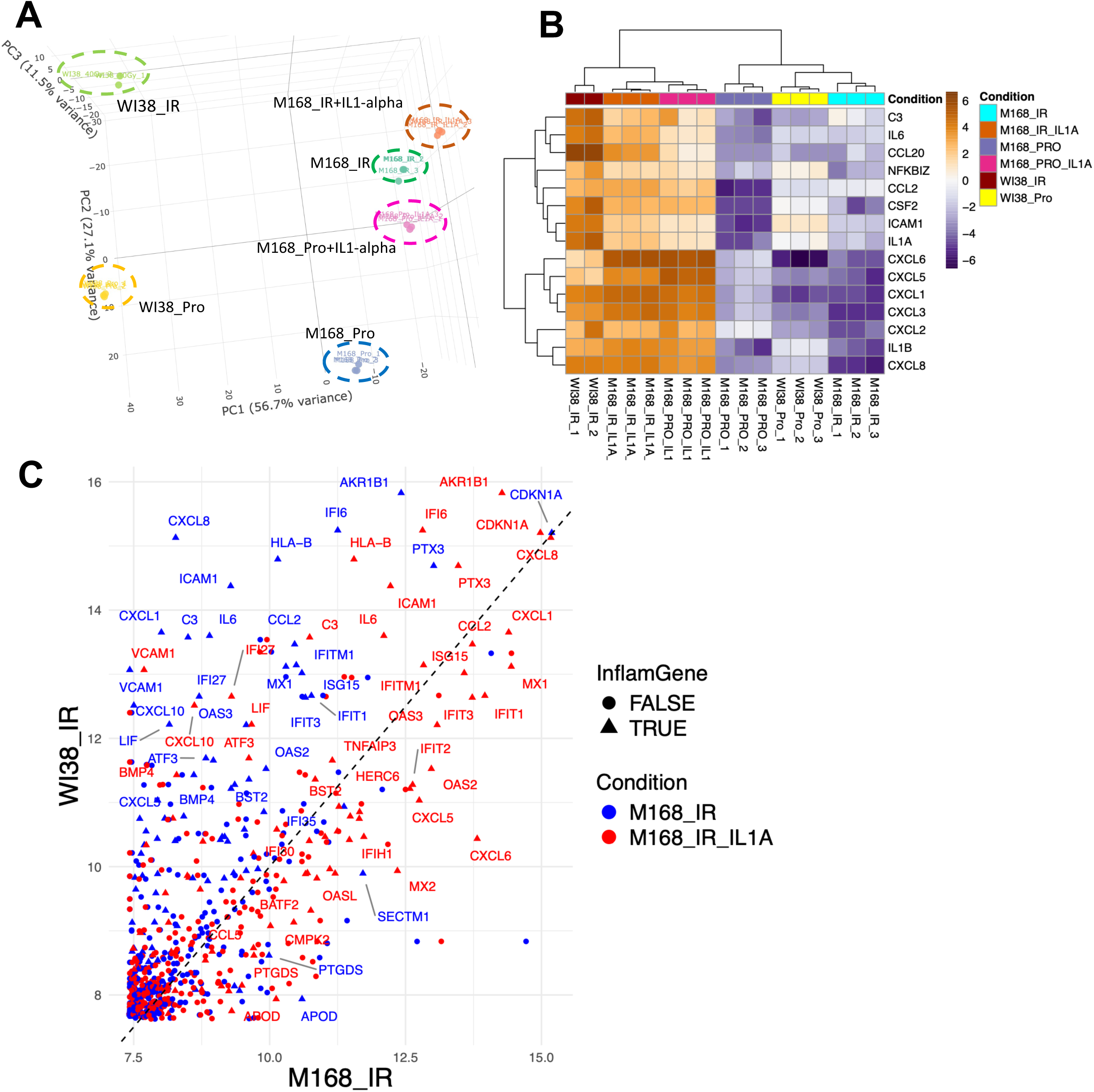
Global transcriptomic effects of the addition for 24 hours of 20 pg/ml IL1-alpha to M168 cells in proliferation or 9 days after 40 Gy X-irradiation. (**A**) 3-dimensional PCA showing separation of WI38 from M168 in PC1, proliferation versus IR or IL1-alpha in PC2, and IR + IL1-alpha from IR in PC3. (**B**) Gene expression heat map of 15 RELA-target genes with high IR-inducible expression in WI38 cells but not M168 cells. Addition of IL1-alpha in proliferation or IR leads to high-level induction of these genes in M168 cells. (**C**) Scatter plot of inflammatory gene expression in WI38_IR cells versus M168_IR cells in the absence (blue dots) or presence (red dots) of IL1-alpha treatment of the M168_IR cells.

We used heat maps of inflammatory gene expression to visualize the effect of IL1-alpha treatment of M168 cells in proliferation and in IR-SEN versus WI38 in proliferation and IR-SEN. From a list of 1110 RELA-target genes, we selected 185 that were highly expressed in either WI38 or M168 cells in IR-induced senescence. We then further selected 55 genes showing the highest variation in all conditions (WI38, M168, proliferation, IR-SEN). The resulting heat map revealed 5 expression clusters (**Fig S12**): cluster-1 contained genes with high expression in WI38 cells relative to M168 cells in both proliferation and IR-SEN, and a weak response to IL1-alpha. Conversely, cluster-2 contained genes with high expression in M168 cells in all conditions and low expression in WI38. Cluster-3 contained genes with low expression in WI38 and M168 cells in proliferation, and induction in both WI38 and M168 after irradiation or IL1-alpha treatment. Cluster-4 contained 3 genes that were induced after IR or IL1-alpha treatment in M168, but not WI38. Finally, cluster-5 was composed of genes with low expression in WI38 and M168 cells in proliferation and that were induced in WI38 cells in IR-SEN, but not M168 cells in IR-SEN. Strikingly, all of these genes were strongly induced in both proliferating and irradiated M168 cells after IL1-alpha treatment. Cluster-5 contained many of the major RELA-induced inflammatory genes of WI38 in IR-SEN, and these were all poorly expressed in M168 cells in proliferation or IR-SEN, but highly induced by IL1-alpha treatment (**Fig. 5B)**.

From a list of 224 interferon-stimulated genes (ISGs), we also selected 90 that showed the highest expression in WI38 or M168 cells in IR-SEN (**Fig S13**). As noted above for the transcriptomic analysis of adult fibroblasts, most ISGs were expressed at lower levels in M168 compared to WI38. As for the RELA-target genes, IL1-alpha treatment increased the expression of many ISGs in proliferating M168 cells, but a synergistic effect was observed for IL1-alpha treatment of M168 cells in IR-SEN.

To more graphically illustrate the effect of IL1-alpha treatment on RELA and IFN target genes, we overlaid scatter plots of gene expression values for WI38 cells and M168 cells in IR-SEN, with or without IL1-alpha treatment of the M168 cells (**Fig 5C**). The scatter plot shows that most of the inflammatory genes are skewed to higher expression in WI38 cells compared to M168 cells in IR-SEN (blue dots), but treatment of irradiated M168 cells with 1L1-alpha generally shifted expression of inflammatory genes towards the M168 cells. These data are consistent with poor inducibility of IL1A and IL1B of M168 cells in IR-SEN explaining the overall poor expression of inflammatory genes in these cells relative to WI38 cells.

### Mechanism of weak DNA-damage inducibility of IL1 in adult fibroblasts

The weak expression of inflammatory genes in adult versus fetal/neonatal fibroblasts appears linked to the weak basal expression of IL1 in these cells. We considered that differences in the expression of trans factors and epigenetic differences in cis-acting regulatory regions govern expression of IL1.

### Epigenetic differences in candidate IL1 enhancers contributing to inflammatory gene expression differences in fetal/neonatal versus adult fibroblasts

The H3-K27me3 repressive mark catalyzed by Polycomb complexes is implicated in cell-type specific gene repression, and one study suggested a contribution of H3-K27me3 repressive marks to the inhibition of inflammatory gene expression in proliferating fibroblasts [25]. We mapped H3-K27me3 genome-wide by Cut & Tag in WI38 fetal lung and adult abdominal (Ab1), breast (M168), and lung fibroblasts (S31). H3-K27me3 was mainly associated with gene bodies and their proximal regulatory sequences in all 4 fibroblasts (75% of features) with the remainder in distal intergenic sequences (**Fig S14A**). Comparing the top 250 genes that are differentially expressed in WI38 fetal fibroblasts and M168 breast fibroblasts, we found an overall inverse correlation with H3-K27me3 signal density and cell-type specific gene expression as expected (**Fig S14B,C**). We next examined specific genomic regions in a genome browser to gauge H3-K27me3 density at RELA inflammatory genes and some candidate regulatory factors. The CDKN2A gene is a well-known target of Polycomb repression in proliferating fibroblasts [25–27]. Compared to CDKN2A, the 72 kb IL1A-1L1B cluster on chromosome 2, the CXCL8 gene on chromosome 4, and the IL6 gene on chromosome 7 showed sparse and low levels of H3-K27me3 signal (**Fig S15**). These data suggest that H3-K27me3 does not play a major role in proximally repressing these inflammatory genes in proliferating fibroblasts, but we cannot rule out a possible repression by H3-K37me3 at unidentified distal regulatory sequences. In contrast to the above, we observed significant H3-K27me3 density at the CXCL1 and CXCL6 genes linked on chr 4 in WI38 fetal lung cells, but not in the adult abdominal Ab1, adult breast M168, or adult lung S31 cells (**Fig S16**). H3-K27me3 at CXCL1 and CXCL6 decreased in senescent WI38 cells in which both genes are expressed. Thus, H3-K27me3 may contribute to repressing CXCL1 and CXCL6 in proliferating WI38 cells, but did not appear to do so in adult fibroblasts. Conversely, ICAM1 and the CXCL10-CXCL11 cluster contained significantly higher levels of H3-K27me3 in the three adult fibroblasts compared to WI38, so it is possible that Polycomb represses these genes to greater extents in adult fibroblasts compared to WI38 (**Fig S16**). In summary, H3-K27me3 levels show significant gene-specific and cell-type specific variability in fibroblasts. Although H3-K27me3 may contribute to the proximal repression of some inflammatory genes in adult fibroblasts, such as ICAM1 and the CXCL10-CXCL11 cluster, it does not seem to explain increased repression of the IL1A and IL1B genes in adult versus fetal fibroblasts.

### Candidate IL1A/IL1B cis-Regulatory Elements with lower activation marks in adult fibroblasts

The IL1A and IL1B genes are tandemly oriented on a 65 kb region of chromosome 2. We reasoned that the low basal expression of IL1A and IL1B in adult versus fetal WI38 fibroblasts might be manifested as cis-regulatory regions with lower accessibility (ATAC-seq) and activation marks (H3K27Ac) in the adult versus fetal WI38 fibroblasts. Global analysis indicated that ATAC-seq and H3K27Ac were enriched in promoters, introns, and distal intergenic sequences relative to exons (**Fig S17**). ATAC-seq and H3K27Ac signals at TSS sites were positively correlated with the expression of genes in WI38 and M168 fibroblasts We then sought peaks with lower ATAC-seq accessibility and/or lower H3K27Ac levels in the adult versus fetal WI38 fibroblasts over a 1 Mb region centered on the 1L1A and 1L1B genes. Two regions within the 35 kb intergenic region separating 1L1A and IL1B fit these criteria.This intergenic region contains a series of candidate cis-regulatory regions according to the Encode consortium [28]. In WI38 fibroblasts, 3 regions had prominent ATAC-seq and H3K27Ac peaks that we labeled candidate IL1-Enh1, Enh2, and Enh3 (**Fig 6**). The Enh2 and Enh3 regions showed markedly lower ATAC-seq and/or H3-K27Ac peaks in the adult versus WI38 fibroblasts in both proliferating, control cells and in senescence induced by ionizing irradiation (**Fig 6**). Addition of IL1-alpha to irradiated M168 cells increased the ATAC-seq and H3-K27Ac peaks in correlation with our previous results showing that IL1-alpha addition activates IL1A and IL1B expression in M168 cells. Furthermore, published ChIP-seq data showed that these candidate enhancers are bound by FOS, CEPB, RELA, and FOXF1 which have all been implicated in inflammatory gene expression in senescence [29–33] (**Fig 6B**). We then targeted the candidate IL1-Enh2 and Enh3 for deletion in WI38 cells to test their importance in inflammatory gene expression.

**Fig. 6.**
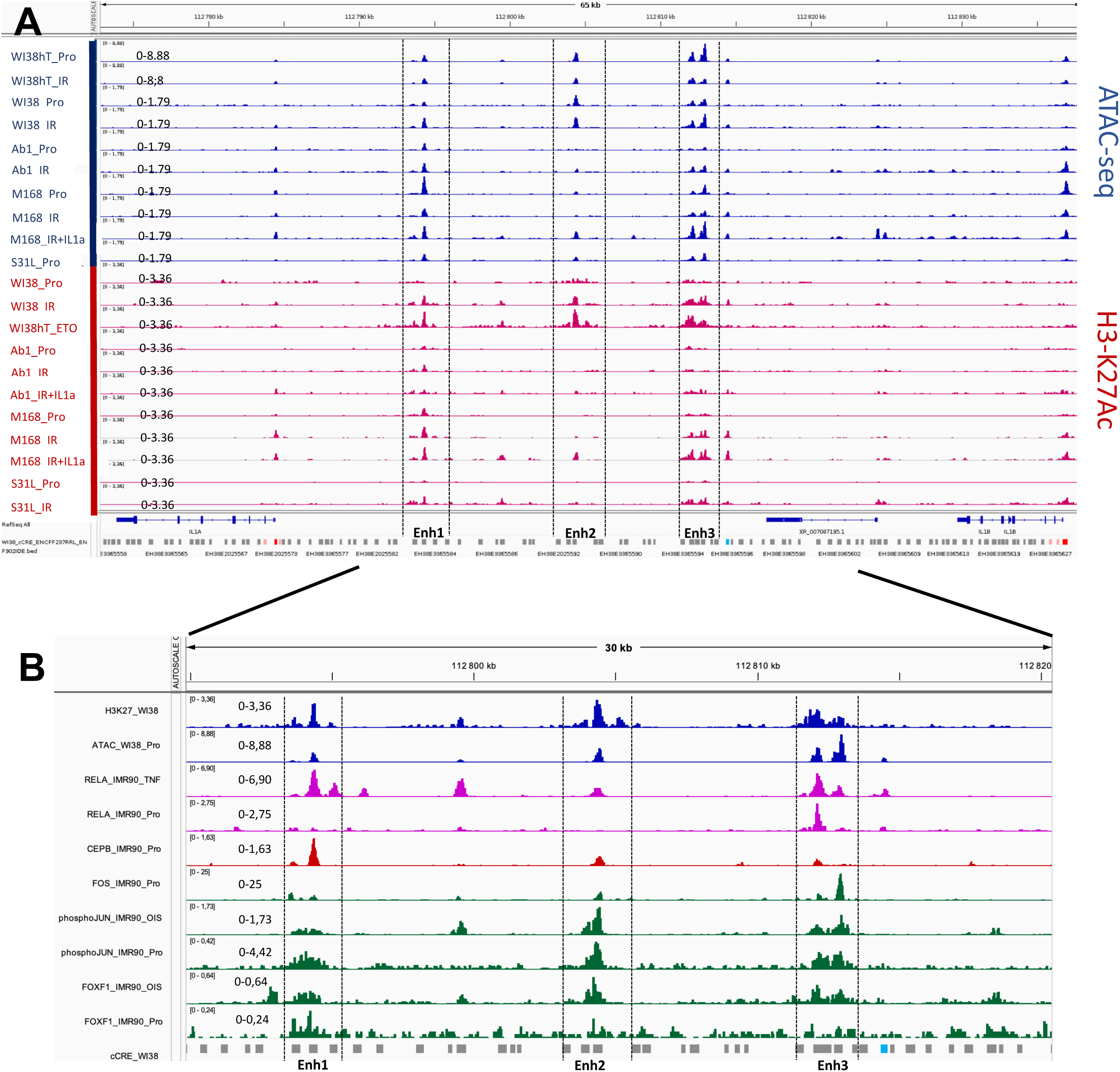
Two candidate enhancers within the IL1 intergenic region show lower accessibility and/or H3-K27Ac in adult versus WI38 fibroblasts and are occupied by key transcription factors implicated in inflammatory gene activation. (**A**) Genome browser screenshots of ATAC-seq (blue) and H3-K27Ac (red) peaks in a 65 kb region IL1A-IL1B. Three prominent peak regions in WI38 fibroblasts representing candidate enhancers are labeled Enh1, Enh2, Enh3. Enh2 and Enh3 show lower ATAC-seq and/or H3-K27Ac signals in adult fibroblasts compared to fetal/neonatal fibroblasts whereas Enh1 shows more comparable signals in all the fibroblasts. Pro = proliferation, IR = 40 Gy X-rays + 10 days, IR + IL1a= 40 Gy + 9 days + 20 pg/ml IL1-alpha for 24h. Also shown are Ref Seq annotation and Encode candidate Cis-acting Regulatory Elements (cCRE). The blue CRE represents a CTCF binding site, red cCREs have TSS promoter-like signatures, pink cCREs have high DNase-accessibility and high-H3K4me3 scores outside of the TSS. (**B**) ChIP-seq and Cut & Run data in IMR90 fetal lung fibroblasts for RELA, CEBPB, FOS, phospho-Jun, and FOXF1 in proliferating cells (Pro) or RELA in IMR90 cels treated with TNF-alpha (TNF), and phosho-JUN and FOXF1 in IMR90 cell in oncogene (RASv12)-induced senescence (OIS). Each sample is auto-scaled to highlight relative peak densities. Note the significant occupancy of RELA at Enh3 even in proliferating cells in which the vast majority of RELA is normally cytosolic. Shown also are the WI38 ATAC-seq and H3-K27Ac peaks to identify Enh1,2,3. The blue cCRE box shows a CTCF binding site.

### Crispr-Cas9 deletion of IL1-Enh2 and Enh3

We anticipated several factors in deciding how to test the functional importance of IL1-Enh2 and Enh3 in the induction of inflammatory gene expression. Since small amounts of exogenous IL1-alpha or IL1-beta can induce the expression of inflammatory genes in our adult fibroblasts, it seemed essential to isolate clones that were homogeneously deleted for the enhancer sequences since even a small fraction of cells that had escaped deletion might express sufficient levels of IL1-alpha and IL1-beta to induce inflammatory gene expression in deleted cells by paracrine activation. Since cloning and amplification of fibroblasts might exhaust their replicative potential, we used an immortalized WI38hTERT cell line to isolate IL1 enhancer deletions. WI38 cells are derived from fetal lung. Although these cells are homogeneously fibroblastic, single-cell RNA sequencing studies have shown that fibroblasts can be classified into numerous subtypes. It was thus important to determine whether all WI38 cells have the potential to induce inflammatory gene expression in senescence induced by IR. Analysis of published single-cell RNA-seq data for proliferating WI38 cells [34] indicated a limited heterogeneity in single-cell gene expression that mainly represented repartition in the cell-cycle phases (**Fig S18A,B**). Comparison of marker genes expressed in 11 different classes of lung fibroblasts [35] indicated that WI38 fibroblasts seem most similar to a fibrosis-associated cluster of fibroblasts with moderate to high expression of CTHRC1, POSTN, FST, and TNC genes, although with low expression of SPP1 that is also expressed in cluster 8 (**Fig S18C**). Thus, WI38 fibroblasts most resemble fibroblasts associated with wound healing involving proliferation, migration, and high ECM production. These results indicated that the WI38 cells were relatively homogeneous as a cell type. To directly test the ability of WI38hTERT clones to induce inflammatory gene expression in IR-induced senescence, we isolated 8 independent clones, induced their senescence, and tested for induction of CXCL8 expression by immunofluorescence. All 8 clones induced expression of CXCL8 in IR-induced senescence (**Fig S19**). We thus concluded that CRISPR/Cas9 deletion of the candidate IL1 Enh2 and Enh3 sequences in WI38hTERT cells would allow us to test their functional significance.

### Crispr/Cas9 deletion of IL1-Enh2 and IL1-Enh3

We created a WI38hTERT cell line constitutively expressing DDCas9, a proteolytically unstable form of Cas9 that can be stabilized by the addition of the small molecule Shield-1 to cells [36]. To delete candidate IL1 enhancers, we added 200 nM Shield-1 to cells for 24 hours to allow accumulation of DDCas9 and we then transiently transfected with synthetic guide RNAs to target DDCas9 to sites surrounding the enhancers (**Fig 7A**). We obtained 2 independent clones deleted for the candidate enhancers IL1-Enh2 and IL1-Enh3 (**Fig S20**). CXCL8 was not induced in these IL1-EnhΔ cells in response to IR, but CXCL8 expression was strongly induced by the addition of exogenous IL1-beta (**Fig. S21**). RT-qPCR experiments showed that both enhancer deletions blocked the IR-induction of 10/10 RELA-target genes that we tested (**Fig 7B**). We conclude that these enhancers are required for the induction of the prime RELA-target genes during IR-induced senescence, but not the acute activation of CXCL8 in response to the strong stimulus provided by the addition of high levels of exogenous IL1-beta.

**Fig. 7.**
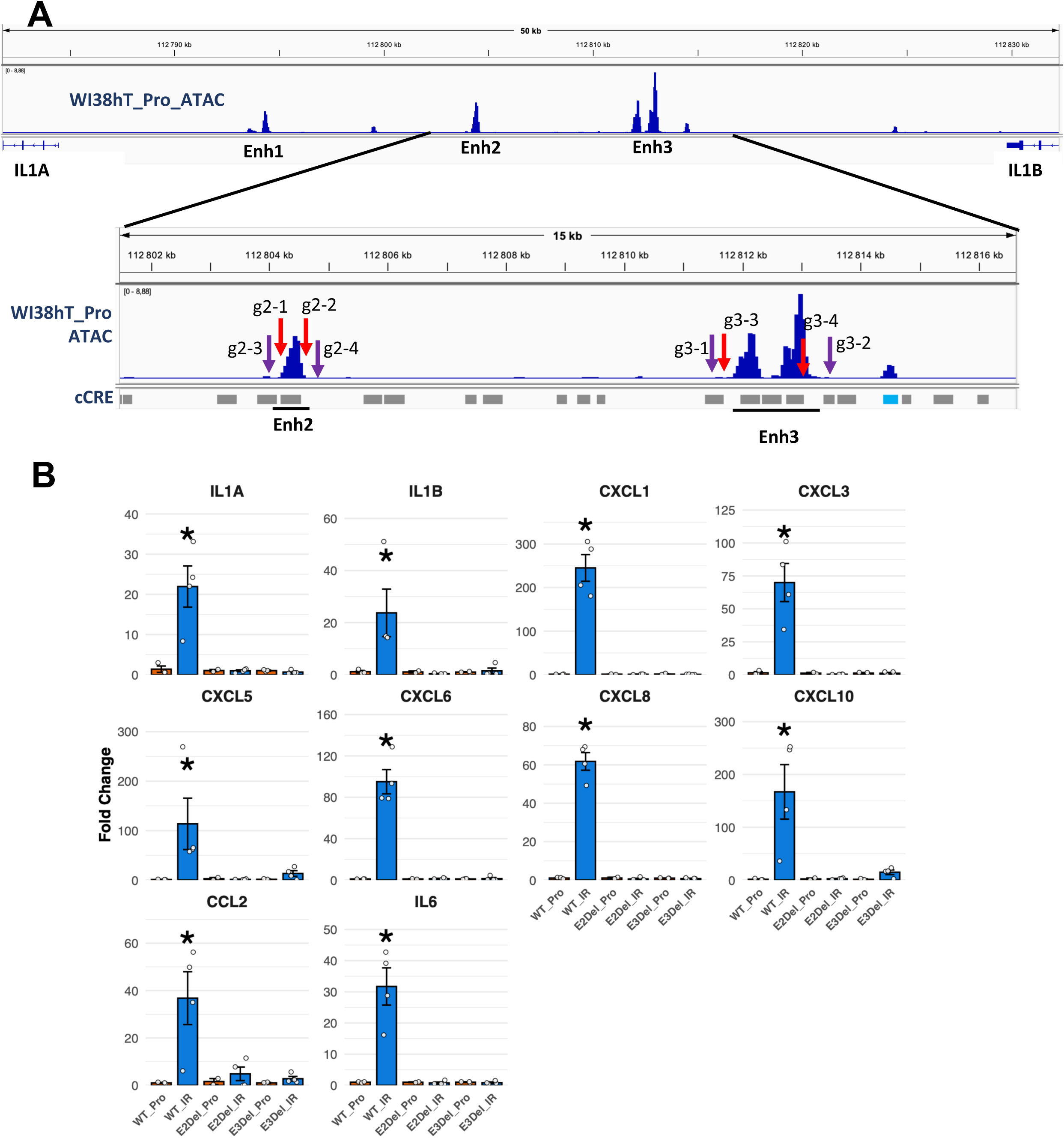
Deletion of IL1-Enh2 or IL1-Enh3 blocks the induction of RELA-target genes in IR-SEN. (**A**) Schema showing the position of pairs of synthetic guide RNAs targeting the deletion of IL1-Enh2 and IL1-Enh3. (B) RT-qPCR of RNA levels for the indicated inflammatory genes comparing the induction level of cells in IR-SEN compared to the same cells in proliferation. Shown are the data points of independent biological replicates for WT cells and for 2 biological replicates of 2 independent Crispr/Cas9 deletions of Enh2 (E2Del) and Enh3 (E3Del). The asterisk indicates that the fold change for WT_IR versus WT_Pro is significantly different (p < .05) compared to the IR versus Pro fold changes for E2Del and E3Del ( mean +/-SEM, Welch’s t-test, n = 4).

### Trans-acting factors potentially contributing to inflammatory gene expression differences in fetal/neonatal versus adult fibroblasts

The differential accessibility of IL1 enhancers in fetal/neonatal versus adult fibroblasts is presumably dependent on the differential activity of trans-acting regulators. One mechanism of differential activation is through differential expression of transcription factors. Our comparative transcriptome analysis revealed some genes that are more highly expressed in fetal/neonatal fibroblasts that may promote inflammatory gene activation in senescence, and some genes that are more highly expressed in adult fibroblasts that may inhibit inflammatory gene expression in senescence. For this analysis, we grouped 3 different fetal lung fibroblasts (WI38, IMR90, MRC5) and 2 different neonatal foreskin fibroblasts (HCA2 and BJ), and we compared them to adult fibroblasts from diverse tissues. Multidimensional scaling (MDS) showed some separation of fetal/neonatal versus adult fibroblasts (**Fig S22A**). With an FDR of 0.01, 482 genes were found to be up-regulated in fetal/neonatal fibroblasts relative to adult fibroblasts and 531 genes down-regulated (**Table S4**). Focusing on transcription factor (TF) genes as candidate regulators, we found 46 TF genes that are expressed at higher levels in fetal/neonatal fibroblasts relative to adult fibroblasts and 49 TF genes that are expressed at higher levels in the adult fibroblasts (**Fig S22B**).

The TWIST2 transcription factor is expressed at higher levels in adult fibroblasts (**Fig S22B**) and TWIST was described as an inhibitor or a co-activator of RELA depending on the context [37–39], so it may conceivably have an anti-inflammatory effect in adult fibroblasts. TF genes expressed at higher levels in fetal/neonatal versus adult fibroblasts may contribute to higher inflammatory gene expression in these cells. The PITX1 and SOX11 transcription factor genes showed the highest differential expression in fetal/neonatal versus adult fibroblasts (**Table S4**). We further confirmed an age-dependent decline in PITX1 and SOX11 expression by analyzing an RNA-seq data collection obtained from dermal fibroblasts of individuals aged 1-98 years [40] (**Fig S23A,B**). Further work is required to test a possible role for these transcription factors in promoting inflammatory gene expression in young versus adult fibroblasts induced into senescence by IR.

We next used ATAC-seq and TOBIAS transcription factor footprinting analyses [41] to identify transcription factor binding motifs that are differentially occupied in proliferating WI38 versus M168 cells (**Fig S23C**). This analysis detected 48 motifs that were bound at higher levels in WI38 versus M168 cells and 46 motifs that were conversely bound at higher levels in M168 cells versus WI38. Interestingly, the TWIST, EN1, PRRX2, and NFIX motifs are predicted to be bound at higher levels in M168 cells and the genes encoding these TFs are indeed expressed at higher levels in M168 cells versus WI38 cells (**Fig S23D**). Further work is required to see if they have anti-inflammatory activity. Notably, a cluster of FOX TFs is predicted to be bound at higher levels in WI38 cells versus M168 cells and our transcriptomics data indicated that FOXF1 and FOXL1 are expressed at higher levels in fetal/neonatal versus adult fibroblasts (**Fig S23E**), and FOXL1 was previously implicated in inflammatory gene expression in lung fibroblasts [42]. FOXF1 showed the highest expression in WI38 cells and had previously been implicated in oncogene-induced senescence (OIS) in fibroblasts [30,43]. We thus decided to test a possible role for FOXF1 in inflammatory gene expression in IR-induced senescence. We obtained 2 different Crispr/Cas9-mediated knock-out mutants of FOXF1 in WI38hTERT cells (**Fig S24**). RNA-seq analysis (**Table S5**) showed a clear separation of WI38 from FOXF1Δ mutants and IR versus proliferation by PCA (**Fig 8A**). GSEA of irradiated FOXF1Δ mutants versus WI38 indicated a down-regulation of both NF-kB (RELA) and Interferon gene sets in irradiated FOXF1Δ versus WI38, and an up-regulation of the Oxidative Phosphorylation gene set (**Fig 8B)**. Heat maps of the major RELA-target genes (**Fig 8C**) and Interferon-Stimulated Genes (**Fig 8D**) confirmed the strong inhibition of inflammatory gene expression in irradiated FOXF1Δ cells. These results are consistent with very recent work showing that siRNA-mediated knockdown of FOXF1 and/or FOXF2 inhibit inflammatory gene expression during OIS of IMR90 fetal lung fibroblasts and DNA-damaged induced senescence of TIG3 fetal lung fibroblasts [31]. It thus seems possible that the increased expression of FOXF1 and FOXL1 in fetal/neonatal fibroblasts compared to many adult fibroblasts contributes to the enhanced inflammatory gene expression during IR-induced senescence of the fetal/neonatal cells.

**Fig. 8.**
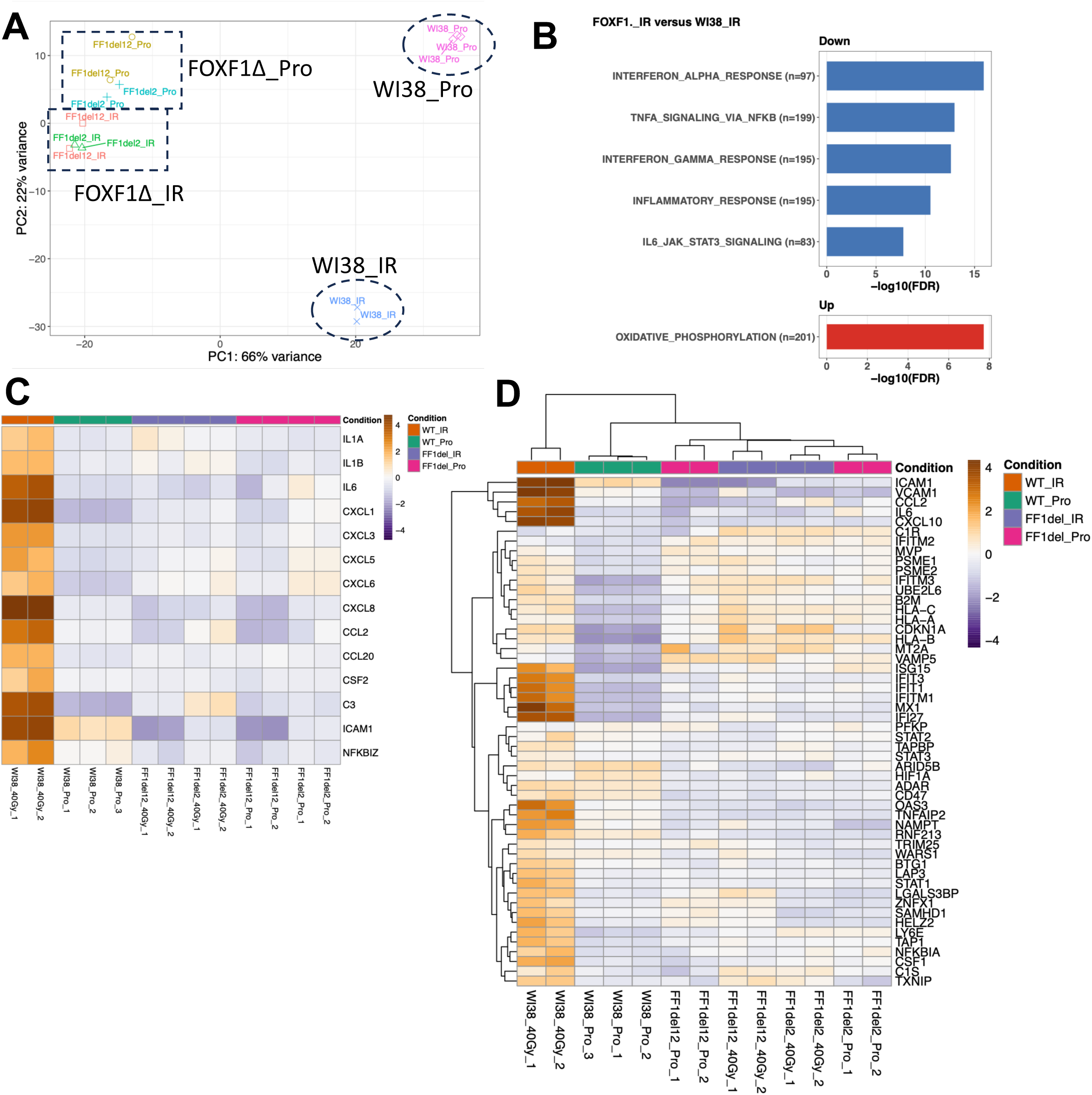
FOXF1-KO mutants of WI38 cells block inflammatory gene expression in IR-SEN. (**A**) PCA showing distinct clustering of WI38 and FOXF1-KO cells in proliferation and IR-SEN. (**B**) Hallmark GSEA shows down-regulation of inflammatory genes in FOXF1 versus WI38 in IR-SEN and up-regulation of the Oxidative Phosphorylation gene set. (**C**) Heat maps of log2FC gene expression for the major RELA-target genes and ISGs (**D**) in IR-SEN for WI38 and FOXF1-KO cells.

## Discussion

The process of cellular senescence has attracted much interest due to its dual beneficial and adverse effects in a context-dependent fashion [1]. The beneficial effects include roles in development, wound healing, and in preventing tumor initiation. The adverse effects are attributed to accumulation of senescent cells during aging associated with stem cell depletion and the chronic secretion of inflammatory factors [4]. However, it is difficult to definitively identify senescent cells in vivo [44,45]. Despite intensive efforts, no universal markers of cellular senescence have been identified. By definition, senescent cells are non-proliferative, but this criterion is not very useful in vivo where most cells are quiescent or post-mitotic. In practice, it is proposed that senescent cells may be identified through phenotypes associated with cell stress and with the increased expression of a subset of genes that are often, but not always, induced in senescence [45]. Transcriptomic markers include the RNAs encoding the cell-cycle inhibitors CDKN1A, CDKN2A, CDKN2B, and a set of RNAs encoding components of the SASP. Of particular interest are RELA-targeted inflammatory genes such as IL1A, IL1B, IL6, CXCL8, and other chemokine genes, since the chronic production of these factors may contribute to aging-related diseases [7]. There are important differences in SASP composition as a function of cell type and stress, but it has been generally accepted that fibroblasts induced into senescence by DNA damage express a high level of these RELA-targeted inflammatory genes [5]. The vast majority of studies describing the transcriptomes of senescent fibroblasts has focused on a small number of fibroblasts derived from fetal lung (WI38, IMR90, MRC5, TIG3) or neonatal foreskin (BJ, HCA2). WI38 fetal lung fibroblasts were developed by Hayflick and were important for vaccine development [46], which motivated the isolation of the other fetal lung fibroblasts. The ready availability of foreskin samples from circumcisions explains their use in deriving fibroblast cell lines [47]. The advantage of studying a small number of well-characterized fibroblast cell lines is that they reduce experimental variability due to genetic and epigenetic variation. The disadvantage is that the results may not be generalizable to the vast number of fibroblast cell types in an adult human. Extensive single-cell sequencing of in vivo fibroblasts have revealed a large transcriptomic diversity dependent on tissue origin, the microenvironment, and pathology [48,49]. The tissue identity of fibroblast transcriptomes is maintained in vitro as are some pathological features. For example, some cancer-associated fibroblasts express inflammatory genes at high levels that are maintained when the fibroblasts are cultivated in vitro [50]. However, some transcriptomic classes defined through single-cell sequencing may depend on the in vivo niche of the cells. Our analysis of a single-cell RNA sequencing data set of proliferating WI38 cells (**Fig S18**) suggested that these fibroblasts express a relatively homogenous transcriptome most similar to that of lung fibroblasts in vivo associated with fibrosis [35]. It is not clear whether fibroblasts of this type result from preferential outgrowth of a subtype that exists in vivo, or whether the in vitro culture with a serum-containing medium induces this type of transcriptome by mimicking fibrotic conditions in vivo. Similar considerations apply to all primary fibroblast cell lines amplified in vitro.

Fetal lung and neonatal foreskin fibroblasts can express both RELA-targeted inflammatory genes and interferon-stimulated genes (ISGs) at high levels during senescence induced by DNA damage or in response to RAF/RAS oncogene expression. Our survey of adult fibroblasts from a diverse set of tissues and individuals indicated that most of them expressed RELA-targeted inflammatory genes at appreciably lower levels compared to the fetal/neonatal fibroblasts during senescence induced by ionizing irradiation (**Fig 4**). About half of these adult fibroblasts induced ISGs at levels similar to irradiated fetal/neonatal fibroblasts, indicating that this branch of inflammatory gene expression dependent on the STAT and IRF transcription factors is regulated independently of the RELA-targeted inflammatory genes [51].

Focusing on an adult mammary fibroblast cell line (M168), we found that these cells also expressed IL6 and CXCL8 at low levels in senescence induced by etoposide, a distinct DNA damaging agent (**Fig S2**). During replicative senescence after long-term passaging in vitro to induced telomere attrition, these cells induced IL6 expression but not IL1A, IL1B, or CXCL1/2/5/6/8, indicating that IL6 can be induced independently of the other RELA-target genes in this condition (**Fig S4**). Finally, the M168 cells did express high-levels of IL1B, IL6, and CXCL8 in response to expression of the B-RAF-V600E oncogene, indicating that adult fibroblasts can induce efficiently RELA-target genes upon hyper-mitogenic stimulation (**Fig S5**).

We noticed that low expression of RELA-targeted inflammatory genes in response to IR was correlated with a low level of nuclear RELA translocation in these cells (**Fig S6**). RELA activation in IR-SEN is dependent on a positive auto-amplification loop involving induction of the IL1A and IL1B genes [22–24]. Interfering with this auto-amplification loop blocks the expression of all RELA-target genes [24]. We found that adult fibroblasts expressed IL1A and IL1B at particularly low levels compared to WI38 in both proliferation and after IR-SEN, suggesting a defect in the initiation of the IL1A and IL1B auto-amplification loop in adult fibroblasts (**Fig 4**). Consistent with this hypothesis, the addition of low levels (20 pg/ml) of either IL1-alpha or IL1-beta was sufficient to strongly activate nuclear translocation of RELA and inflammatory gene expression in both proliferating and irradiated cells (**Fig S10, S11, Fig. 5**). We then sought epigenetic differences that might explain the poor inducibility of IL1A/IL1B in 4 adult fibroblasts versus WI38. We did not observe significant amounts of the Polycomb repressive H3-K27me3 mark in the IL1A/IL1B region of our fibroblasts suggesting that direct Polycomb repression does not explain differences in IL1 inducibility in adult versus WI38 fibroblasts (**Fig S15**). In contrast, mapping of H3-K27Ac and ATAC-seq peaks revealed 2 candidate enhancers within the IL1A/IL1B intergenic region that appeared less accessible and/or with less H3-K27Ac in the adult versus WI38 fibroblasts (**Fig 6**). Remarkably, both candidate enhancers are occupied by AP1 family pioneer transcription factors (FOS, phospho-JUN), FOXF1, CEBPB, and RELA, all of which are necessary for inflammatory gene expression in senescence [29–33,52]. Crispr/Cas9 deletion of either of these putative enhancers in WI38 cells blocked the expression of RELA-target genes in IR-SEN, but these cells still induced inflammatory genes in response to exogenous IL1-beta (**Fig 7, Fig S21**). Our current model to explain these results is that strong signaling by exogenous IL1 or TNF-alpha leads to massive translocation of RELA into the nucleus in an active form that can bind to open enhancers already occupied to some extent by at least AP1, CEBPB, and FOXF1, resulting in rapid activation of IL1A, IL1B and other target genes. In contrast, relatively weak signaling in response to DNA damage leads to nuclear translocation of a small fraction of RELA in our fibroblasts. In our low responding adult fibroblasts, this weak mobilization of nuclear RELA is insufficient to significantly activate IL1A and IL1B transcription, and thus unable to activate the RELA auto-amplification loop. In fetal lung and neonatal foreskin fibroblasts that contain multiple accessible enhancers primed with H3-K27Ac, this level of nuclear RELA is sufficient to allow IL1A and IL1B transcriptional activation in at least a fraction of cells. The resulting production of IL1-alpha and IL1-beta can then trigger auto-amplification of RELA signaling. We note however, that even for the fetal/neonatal fibroblasts, the level of RELA activation in response to IR is insufficient to allow inflammatory gene expression in all cells in the population (**Fig 1, Fig S6**). This conclusion is independently supported by single-cell RNA sequencing of WI38 fibroblasts in senescence induced by DNA damage in which only a fraction of cells in the population expresses RELA-target genes [53]. In contrast, strong RELA activation by exogenous IL1 and TNF-alpha can activate inflammatory gene expression in nearly all cells in the population (**Fig 1, Fig S6**). Consistent with this model, previous work has shown that acute activation of inflammatory gene expression by TNF-alpha occurs without modification of chromatin loops around the IL1A and IL1B genes [54], whereas activation of these genes in response to the RAS oncogene is associated with a new chromatin loop architecture at this locus that may bring together distal enhancers to favor IL1A/IL1B expression [55].

The differential accessibility and H3K27-acetylation of two IL1 enhancers in adult versus fetal/neonatal fibroblasts is presumably linked to the differential expression of some regulatory factors in these fibroblasts. The signaling pathways and transcription factors implicated in senescent inflammatory gene expression are numerous and complex [5]. Differences between fetal/neonatal and adult fibroblasts may involve multiple elements. Our transcriptome analysis revealed some potential anti-inflammatory transcription factors (TWIST) are expressed at higher levels in our adult fibroblasts compared to fetal/neonatal fibroblasts and conversely, some transcription factors (PTX1, SOX11, NFE2L3, NFAT5, HMGA2, FOXF1, FOXL1) are expressed at higher levels in fetal/neonatal fibroblasts and may potentially facilitate inflammatory gene expression. Bioinformatic ATAC-seq transcription factor fingerprint analysis indicated increased occupancy of the FOXF1 DNA binding motif in WI38 cells compared to adult fibroblasts (**Fig S23C**). We thus generated Crispr/Cas9 KO mutants in WI38hTERT cells and found that the inactivation of FOXF1 blocked inflammatory gene expression in these cells upon IR-SEN (**Fig 8**). This result is in agreement with recent work showing that siRNA mediated knock-down of either FOXF1 or FOXF2 blocked inflammatory gene expression of fetal lung fibroblasts induced into senescence by expression of the RAS oncogene or DNA damage [31]. These authors further showed that FOXF1 forms a complex with AP-1 and colocalizes with AP-1 at multiple candidate enhancers for inflammatory genes. The observation that siRNA knock-down of either FOXF1 or FOXF2 could inhibit inflammatory gene expression may indicate that these factors function as heterodimers, as suggested by the BioGRID protein interaction database (https://thebiogrid.org/108583/summary/homo-sapiens/foxf1.html) and predicted by the Predictome database. The increased expression of FOXF1 in fetal/neonatal versus fibroblasts may thus contribute to their greater inflammatory response in senescence compared to adult fibroblasts. FOXF1 shows tissue-specific and cell-type specific expression (https://www.proteinatlas.org/ENSG00000103241-FOXF1/single+cell) with developmental roles in the lung [56]. It is highly expressed in fetal lung fibroblasts, moderately expressed in adult lung, gingiva, palate, vocal fold fibroblasts and neonatal foreskin fibroblasts, and shows low expression in adult fibroblasts from the breast and abdomen (**Fig S23E**). The levels of FOXF1 expression are not perfectly correlated with inflammatory gene expression in that neonatal foreskin fibroblasts express higher levels of inflammatory genes in IR-SEN compared to adult gingiva, palate, and vocal fold fibroblasts for similar levels of FOXF1 expression in these cells. We thus feel that other factors must contribute to differences in inflammatory potential between adult and fetal/neonatal fibroblasts. FOXF1 was previously shown to contribute to the proliferative arrest of BJ foreskin fibroblasts in response to expression of the RAS oncogene [30,43], although it does not appear to bypass proliferative arrest induced by DNA damage or RAS expression in fetal lung fibroblasts [31]. Further studies are required to understand the role of FOXF1 in the OIS proliferative arrest, and whether other transcription factors can substitute for FOXF1 with regards to the proliferative arrest in fibroblasts that lack FOXF1.

In conclusion, we find that many adult fibroblasts express low levels of RELA-targeted inflammatory genes in senescence induced by DNA damage compared to fetal/neonatal fibroblasts, and we trace this difference to the activity of cis-acting enhancers and to the expression of the FOXF1 transcription factor. Nevertheless, since the adult fibroblasts induce these genes in the presence of low-levels of exogenous IL1-alpha, IL1-beta, or TNF-alpha, they will express these genes in vivo in an inflammatory environment in which recruited immune cells secrete these inflammatory triggers.

### Limitations of this study

Our study has numerous limitations. Amongst others, we examined a limited number of adult primary fibroblasts from a limited number of tissue samples. It would be worthwhile extending this survey. Furthermore, we did not assay DNA methylation that might be an additional repressing factor for inflammatory gene expression in some adult fibroblasts.

### Resource Availability

DNA sequences were deposited at the ArrayExpress of the European Bioinformatics Institute under the following accession numbers: E-MTAB-14201 (RNA-Seq of irradiated primary human fibroblasts), E-MTAB-16030 (RNA-seq of mammary M168 fibroblasts, with and without irradiation and with an without IL1-alpha treatment), E-MTAB-14625 (ATAC-seq or primary human fibroblasts), E-MTAB- (Cut & Tag H3-K27ac and H3-K27me3 of primary human fibroblasts), E-MTAB-17208 (RNA-seq of irradiated FOXF1-KO cells).

## Materials and Methods

### Primary cells and culture conditions

Primary fibroblasts were described previously [19] or obtained for this study as indicated in **Table S1** with informed consent from donors. Cells were cultivated in DMEM (4.5 g/l glucose) + 10% fetal bovine serum with stabilized glutamate, pyruvate, and penicillin/streptomycin in a humidified incubator with 5% CO2 and 5% O2 unless stated otherwise. Inx- some experiments, purified recombinant IL1-alpha (Thermo Fisher PHC0011), IL1-beta (Thermo Fisher 200-01B), or TNF-alpha (Thermo Fisher 300-01A), in PBS were added to cells at the indicated concentrations and incubation times. Primary cells were used within 10 population doublings unless stated otherwise. WI38 cells were purchased from the ATCC. WI38hTERT cells were derived from WI38 as described previously [12].

### x-ray-induced senescence

We followed the protocol developed by the Campisi lab to induce senescence and SASP expression by ionizing irradiation [8]. Fibroblasts were grown to confluence in 10 cm dishes to enrich for cells in the G1 phase of the cell cycle and then irradiated with 10, 20, or 40 Gy of 120 kV X-rays at 2 Gy/minute produced by a Faxitron Model 43855D X-ray generator. The next day, cells were split into 3 x 10 cm dishes and incubated for a total of 9 further days with fresh medium applied every 3 days. We assayed for senescence induction by EdU incorporation and SA-ß-galactosidase assays at 10 days post-irradiation. We found that 40 Gy irradiation was necessary to ensure that at least 80% of cells were induced into senescence. We needed to use higher doses in our work compared to previous studies because our irradiator produced lower energy X-rays (120 kV).

### Replicative senescence of M168 mammary fibroblasts

Cells were passaged in an incubator with 5% CO2 in ambient oxygen to accelerate replicative senescence until less than 10% of cells showed incorporation of EdU. This required approximately 80 population doublings.

### M168 senescence induced by B-RAF-V600E

M168 cells were infected with a lentiviral pTRIPz-B-RAF-V600E preparation [57] and transduced cells were selected for resistance to 1 µg/ml puromycin. B-RAF-V600E expression was induced by the addition of 100 ng/ml doxycycline for 4 days before extracting RNA for RT-qPCR analysis of inflammatory gene expression.

### Crispr/Cas9 deletions

A stable cell line expressing DDCas9 was produced by infecting WI38hTERT cells with a DDCas9 lentivirus [36], followed by FACS purification of cells expressing the Venus fluorescent protein. To target genomic deletions, DDCas9 was stabilized by the addition of 200 nM Shield-1 for 24 hours and cells were then transfected with 2 Alt-R sgRNAs (**Table S6**), synthesized by Integrated DNA Technologies, at a final concentration of 40 nM in Lipofectamine RNAiMAX. The 2 sgRNAs defined the borders of a genomic DNA fragment that we targeted for deletion. Clones were isolated by dilution into 10 cm dishes and incubation until isolated clones were visible in a phase contrast microscope. Clones were then recovered in a drop of PBS + trypsin and transferred into wells of a 96-well plate for amplification. Clones containing genomic deletions were identified by PCR after rapid extraction of DNA using the heat denaturation and proteinase K protocol [58] from cells in a 96-well plate. The sequence of PCR primers flanking the IL1Enh2 and IL1Enh3 deletions that were used to identify deletion clones are shown in **Table S6**.

For the IL1EnhΔ clones, we found that most clones contained a deletion of the targeted fragment on one allele, but on the second allele, we found microdeletions at each sgRNA-targeted cut site without deletion of the intervening genomic DNA. We thus designed a second set of 2 sgRNAs to target the interior region within the second allele and we did a second round of transfection, cloning, and PCR screening to identify clones containing a homozygous deletion of the IL1Enh2 and Enh3 as indicated in **Fig 7** and **Fig S20**. For the FOXF1-KOs, we screened clones by immunofluorescence and Western blotting to identify 2 independent knock-out clones (**Fig S24**).

### EdU proliferation assays

Fibroblasts were seeded into 96-well plates at 15000 cells per well. The next day, fresh medium containing 1 µM 5-ethynyl-2′-deoxyuridine (EdU) was added to cells and incubation was continued for 24 hours before fixing the cells with PBS + 3.7% formaldehyde for 10 minutes and processing the cells for detection of EdU in the nucleus with Alexa488-azide or Alexa594-azide as described [59]. We visualized and quantified nuclear fluorescence with a Thermo Fisher CX5 high-content screening platform.

### Immunofluorescent assays

Fibroblasts were seeded into 96-well plates at 15000 cells per well. The next day, cells were fixed with PBS + 3.7% formaldehyde for 10 minutes, permeabilized for 5 minutes with PBS + 0.5% Triton-X-100 and washed with PBS + 0.1% Tween-20. Cells were then incubated with a 1/500 dilution of primary antibodies in PBS + 0.1% Tween-20 for 1 hour at room temperature (RT) followed by washing with PBS + 0.1% Tween-20 and then incubation with a 1/1000 dilution of secondary antibody in PBS + 0.1% Tween-20 for 1 hour at RT followed by washing with PBS + 0.1% Tween-20 and a final incubation with PBS + 0.1 µg/ml DAPI to stain nuclear DNA. The following primary antibodies were used: mouse monoclonal anti-CXCL8 (Thermo Fisher M801), goat anti-IL6 (R & D AF206), rabbit monoclonal anti-RELA (CST-8242), rabbit monoclonal anti-FOXF1 (Abcam ab168383). The secondary antibodies were anti-IgGs directed againt mouse, rabbit, or goat IgG coupled to Alexa-488 or Alexa-594 (Thermo Fisher). We visualized cellular fluorescence with a Thermo Fisher CX5 high-content screening platform. For RELA, quantification was restricted to nuclear fluorescence, whereas CXCL8 and IL6 in the secretory apparatus of the cells was quantified in a cytosolic ring in the cells around the nucleus.

**Senescence-associated beta-galactosidase assays** were performed as described [60].

### Western blots of FOXF1

Approximately 2 million cells were resuspended in 300 µl of a protein extraction buffer containing: 50 mM Tris (pH 7.5), 150 mM NaCl, 1M Triton-X-100, 0.1% SDS, and Roche protease inhibitor cocktail. Resuspended cells were incubated 15 minutes on ice, and then spun 21,000 x g for 15 minutes at 4°C. The supernatant protein extract was transferred to a new tube and the protein concentration was determined with the Pierce 660 nm protein assay reagent. 2.5 µg of protein extract was electrophoresed in SDS-12% polyacrylamide mini-gels and proteins were then transferred to nitrocellulose membranes. Membranes were blocked with PBS + Li-Cor Intercept (1:1), and then incubated with a 1/500 dilution of rabbit monoclonal anti-FOXF1 (Abcam ab168383) in PBS + Intercept + 0.1% Tween-20 for 1 hour at RT. After washing, the membrane was incubated with a 1/5000 dilution of anti-rabbit-800 antibody (Li-Cor) in the same buffer for 1 hour at RT. The fluorescent membrane was then scanned on a Li-Cor Odyssey.

### RNA purification

For cells in 10 cm plates, cells were trypsinized, washed with PBS, and the cell pellet was resuspended in Machery-Nagel Nucleospin RNA kit lysis buffer and RNA was purified from spin columns. For cells in 6-well plates, cells were lysed by scraping directly in lysis buffer and RNA was purified with the Machery-Nagel Nucleospin RNA XS kit. RNA was quantified with a Nanopore spectrometer.

### RT-qPCR

500 ng RNA was reverse-transcribed into cDNA using random primers and Maxima reverse transcriptase (Thermo Fisher). A 1/5 dilution of the cDNA was used for qPCR using the Luminaris Color HiGreen qPCR Master Mix (Thermo Fisher) and an Applied Biosystems CX96 thermocycler. The ΔΔCq method was used to quantify relative fold change of cDNA levels using GAPDH as the reference gene. The primers used for qPCR are specified in **Table S6**.

### RNA-sequencing

PolyA RNA-sequencing libraries were prepared with the NEBNext polyA magnetic isolation module, Ultra II Directional RNA kit, and NEBNext multiplex oligos starting with 1 µg of total RNA.

### ATAC-seq

Tn5 was purified and assembled with oligonucleotides as described [61], and ATAC-sequencing was performed with 50,000 cells using the OMNI-ATAC protocol [62].

### Cut and Tag

ProteinA-Tn5 was purified as described (https://www.protocols.io/view/3xflag-patn5-protein-purification-and-meds-loading-j8nlke4e5l5r/v1), and Cut and Tag was performed with 50,000 cells using the nuclear permeabilization protocol [63] with 1 µg of anti-H3-K27-me3 (Abcam 6002) or 1 µg of anti-H3-K27-Ac (Abcam 4729) antibodies.

### Sequencing

High-throughput sequencing of multiplexed DNA pools was performed by the I2BC NGS platform using Illumina NextSeq 500 or 550 sequencers.

### Bioinformatics

The code used for these analyses has been deposited to Zenodo (10.5281/zenodo.20817526), but we summarize the steps below. Quality control metrics for the ATAC-seq and Cut & Tag data are shown in **Table S7**.

Sequences were demultiplexed with Illumina bcl-convert and adapters were trimmed with Cutadapt [64] by the I2BC NGS platform. The trimmed fastq sequences were quality controlled with FastQC.

### RNA-seq bioinformatics

Fastq sequences were mapped to the hg38 transcriptome using salmon (v 1.10.3) with the salmon_partial_sa_index and validateMappings and gcBias options [65]. The resulting quant files were then imported into an annotated summarized experiment with tximeta [66] and the counts from transcripts were aggregated to the gene level. The counts data were variance-stabilize transformed for PCA analysis. Differential-gene expression analysis was performed with limma-voom [67]. Gene set enrichment analysis was done with camera [68]. For some RNA-seq data, the fastq sequences were aligned to the hg38 genome with Star (v2.7.11b) to output bam files that were indexed with samtools and then converted to bigwig files using Deeptools (v3.5.6) with CPM normalization [69].

### ATAC-seq bioinformatics

Fastq reads were mapped to the hg38 genome with Bowtie2 using the parameters --end-to-end, --very-sensitive, --no-mixed, --no-discordant, -I 25, -k 1, -X 1000. The sam output was converted to bam, indexed, duplicates and mitochondrial sequences were removed, and quality control was verified with flagstat and idxstats using samtools and sambamba [70]. Deeptools was used to convert the bam files to bigwig with CPM normalization whilst removing blacklisted regions within the Encode GRCh38_ExclusionList2020_ENCFF356LFX.bed file. Quality control of the ATAC-seq data was evaluated with ataqv [71]. Differential transcription factor fingerprinting and occupancy predictions were performed with the TOBIAS snakemake pipeline [41].

### Cut & Tag bioinformatics

Alignment for H3-K27Ac and H3-K27me3 fastq reads to the hg38 genome was as for ATAC-seq above. Bigwig conversion used CPM normalization for H3-K27Ac, but RPGC (reads per genomic content) for the broad H3-K27me3 marks.

### Peak calling

We used macs3 callpeak with parameters –call-summits -q .01, -B –SPMR for narrow peak H3-K27Ac and ATAC-seq bam files, and parameters –broad, --broad-cutoff 0.05,--nomodel, for the broad peak H3-K27me3 bam files.

## Supporting information

TableS1_Primary Fibroblasts

TableS2_FbIR_DEG,TPM

TableS3_M168_IL1-alpha_DEG, TPM

TableS4_FetalNeonatalvsAdult_DEG,TPM

TableS5_FOXF1-KO_DEG,TPM

TableS6_sgRNAs,qPCR-oligos

TableS7_QC_ATAC,CutTag

## Acknowledgements.

This paper is dedicated to the memory of Reem Hamed who tragically passed away before this manuscript was submitted for publication. Reem was supported by a scholarship from the Université Paris-Saclay International Master’s Program and a doctoral contract with the French Ministry of Research and Higher Education. The present work benefited from the Imagerie-Gif core facility, supported by I’Agence Nationale de la Recherche (FBI ANR-24-INBS-0005 (BIOGEN); SPS ANR-17-EUR-0007, EUR SPS-GSR, and we acknowledge the sequencing and bioinformatics expertise of the I2BC High-throughput sequencing facility, supported by France Génomique (funded by the French National Program “Investissement d’Avenir” ANR-10-INBS-09). The DDCas9 lentivirus was produced by the Genetic Engineering and Expression platform (CIGEx) of the Institut de Biologie François Jacob at the CEA/Fontenay-aux-Roses. We received funding from the CEA/EDF radiobiology program and from LVMH.

