## Supplementary Figures for "Differential Enhancer Activity and FOXF1 Levels Contribute to Higher Inflammatory Gene Expression of Fetal/Neonatal Versus Adult Fibroblasts in IR-induced Senescence"

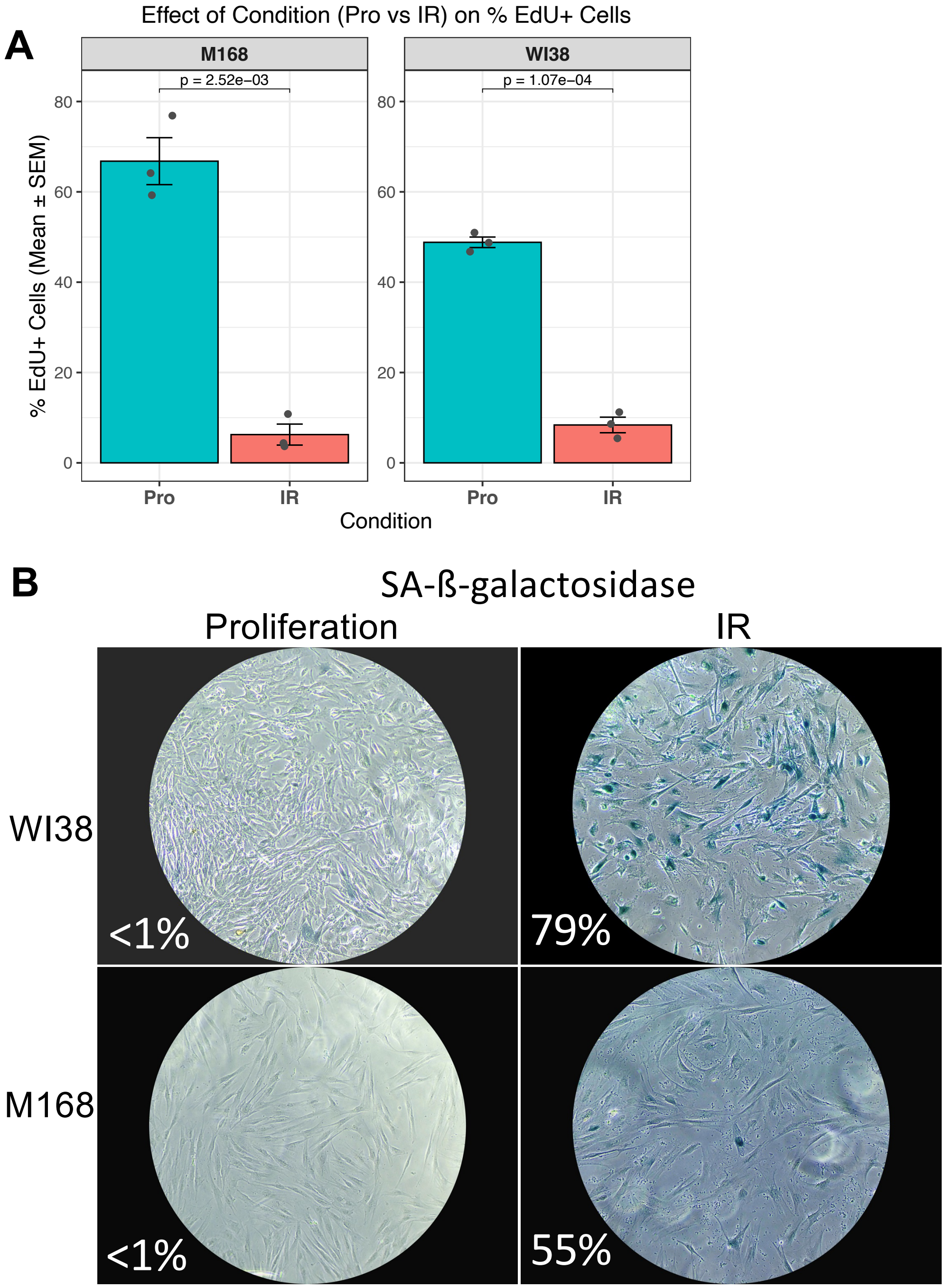


**Fig S1.** Ionizing irradiation (IR) induces senescence of adult mammary M168 and fetal lung WI38 fibroblasts. 10 days post-irradiation (IR), proliferation (Pro) was assayed by (**A**) EdU incorporation (mean +/- SEM, n=3, Welch's t-test) and (**B**) by the senescence-associated ß-galactosidase assay.

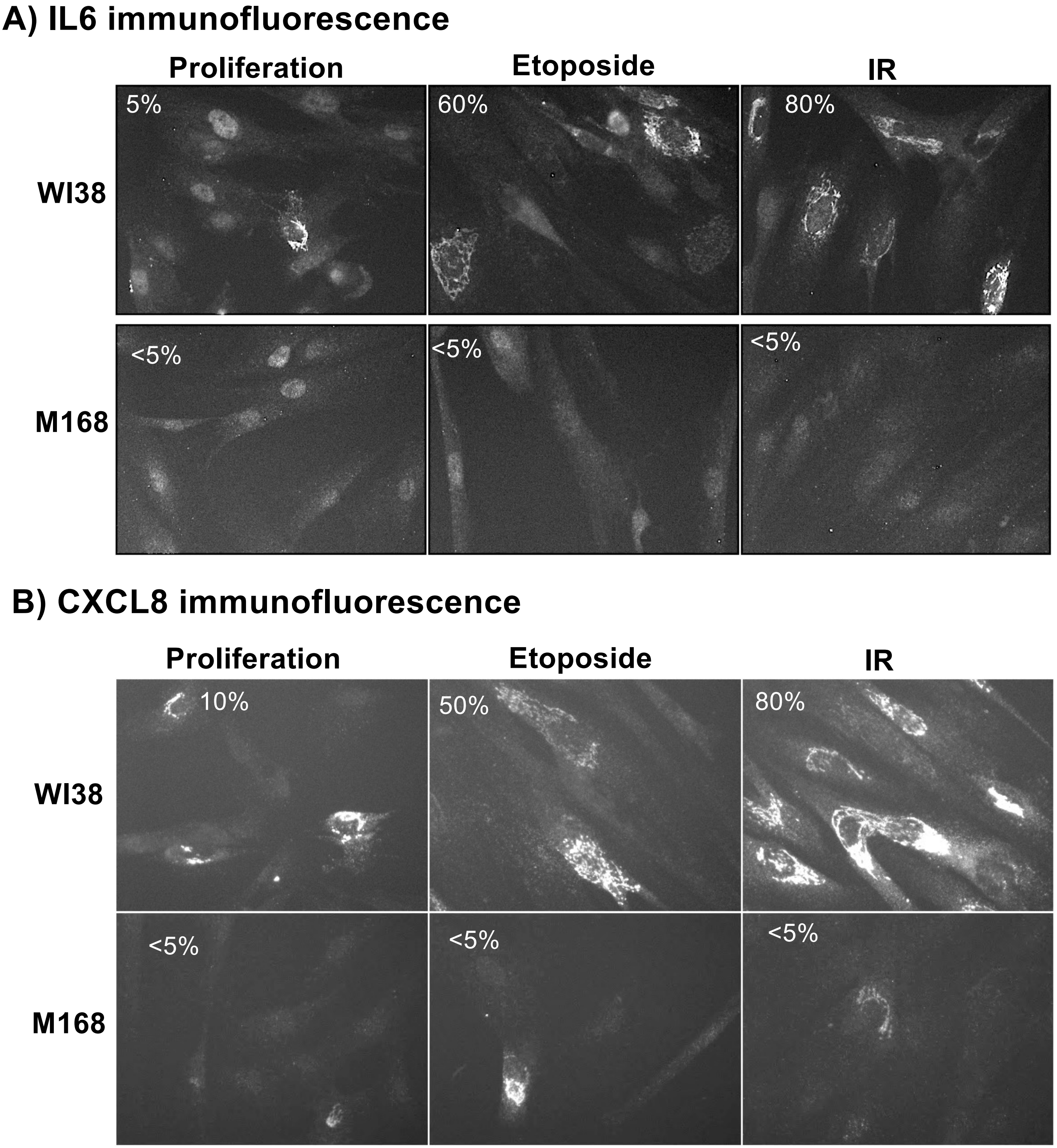


**Fig S2.** IL-6 (**A**) and CXCL8 (**B**) immunofluorescence showing that M168 fibroblasts express low levels of these inflammatory factors in senescence induced by etoposide or IR relative to WI38 fibroblasts. The percentage of positive cells is shown.

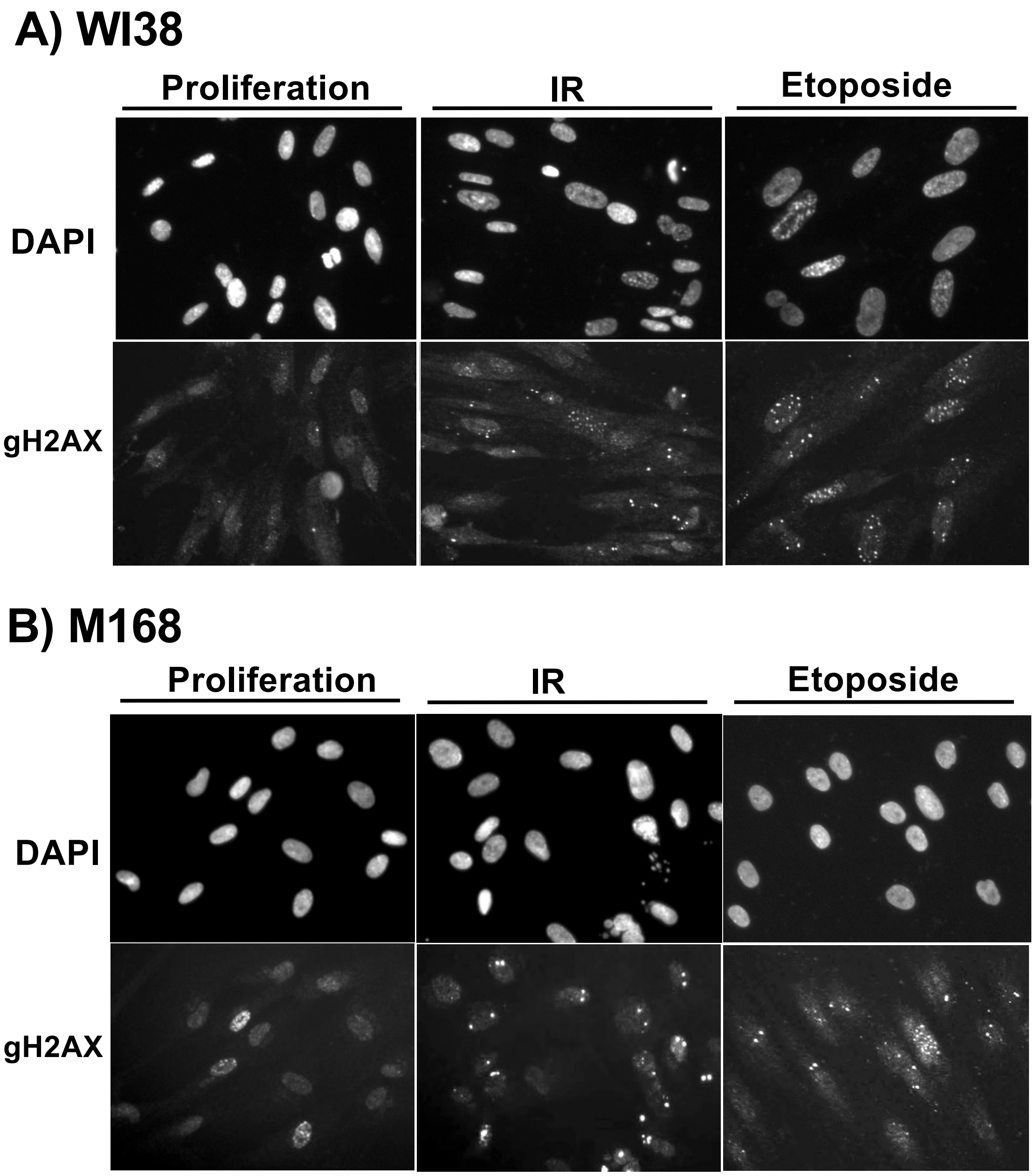


**Fig S3.** gamma-H2AX (gH2AX) immunofluorescence shows that the majority of WI38 (**A**) and M168 (**B**) fibroblasts contain one or more persistent gamma-H2AX foci 10 days after X-irradiation (IR) or treatment with etoposide.

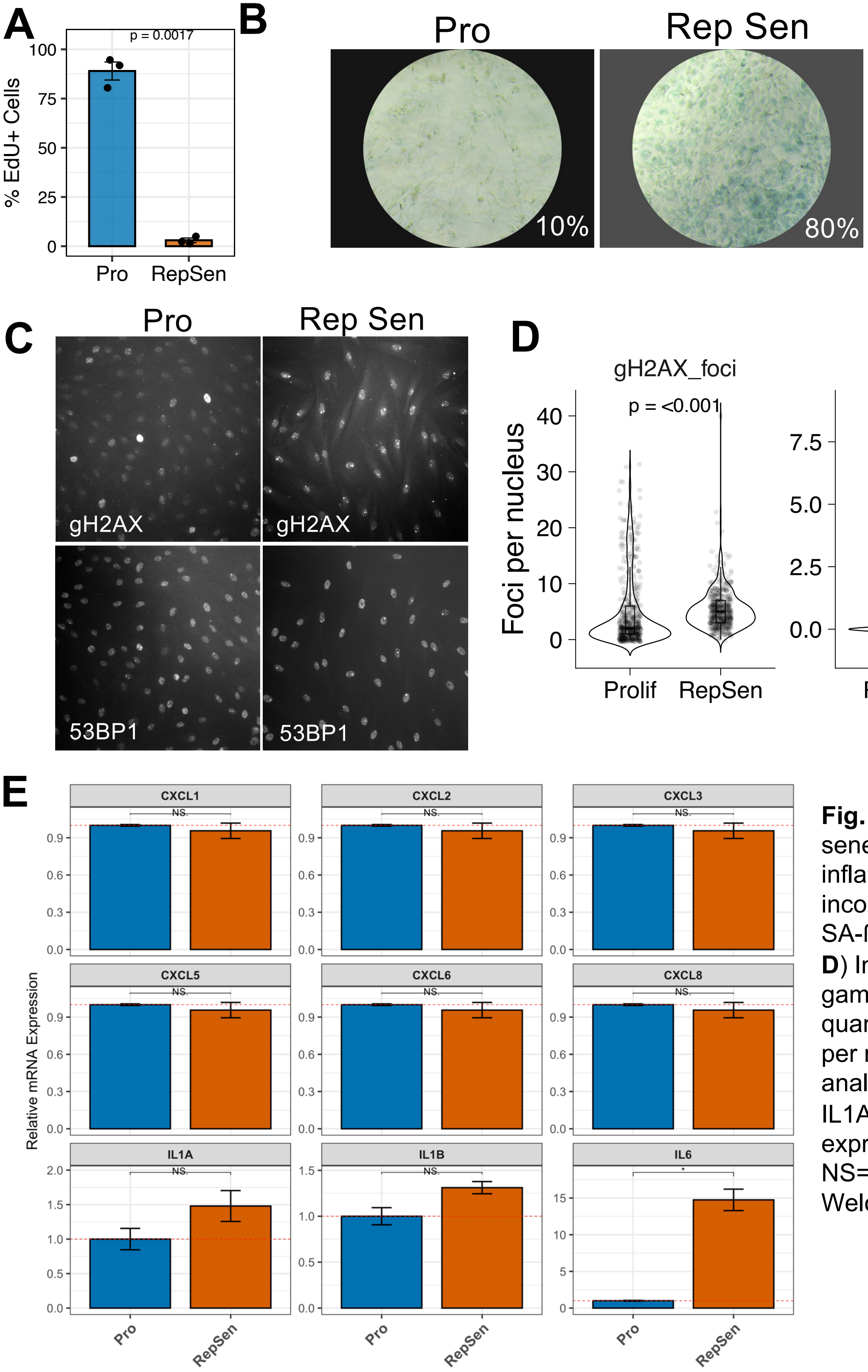


**Fig. S4**. M168 cells in replicative senescence express low levels of inflammatory genes. (**A**) EdU incorporation assays, n=3. (**B**) SA-ß-galactosidase assays. (**C, D**) Immunofluorescent staining for gamma-H2AX and 53BP1 and quantification of the number of foci per nucleus. (**E)** RT-qPCR analysis of CXCL1/2/3/4/5/6/8, IL1A,IL1B, and IL6 gene expression (mean +/- SEM), n=3. NS= not significant, * = p < .05, Welch's t-test.

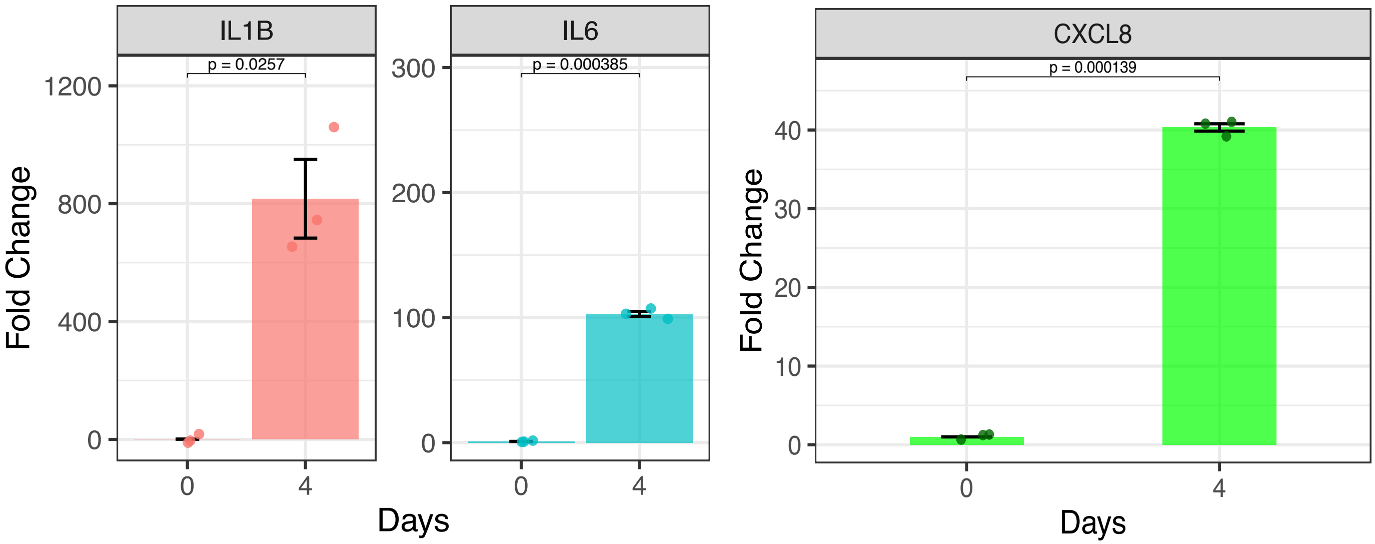


**Fig S5.** RT-qPCR analysis of IL1B, IL6, and CXCL8 gene expression in M168 fibroblasts containing a ptet-ON-B-RAF-V600E construction allowing doxycyline-inducible expression of the B-RAF-V600E oncogene (mean +/- SEM, n=3, Welch's t-test). The fold change gene expression is shown for cells induced for 4 days with 100 ng/ml doxycyline relative to uninduced cells.

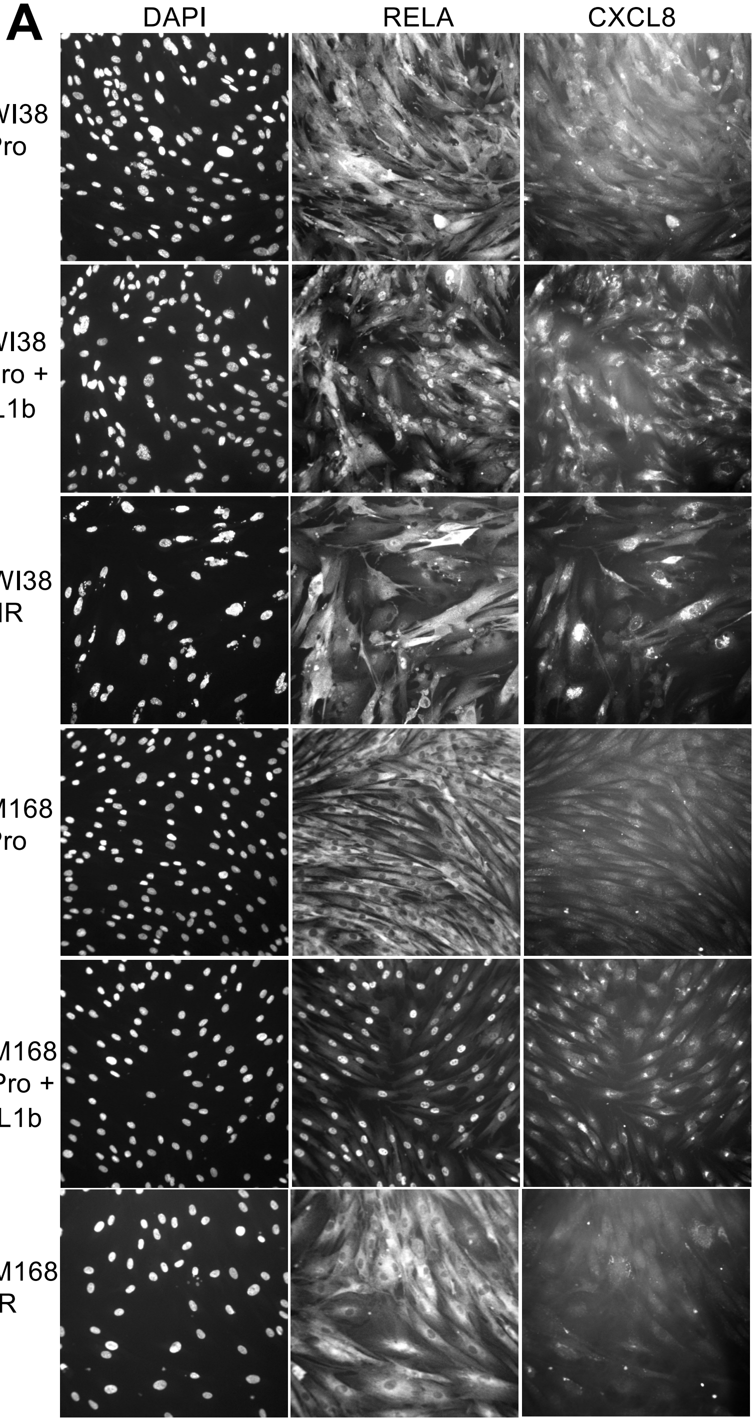


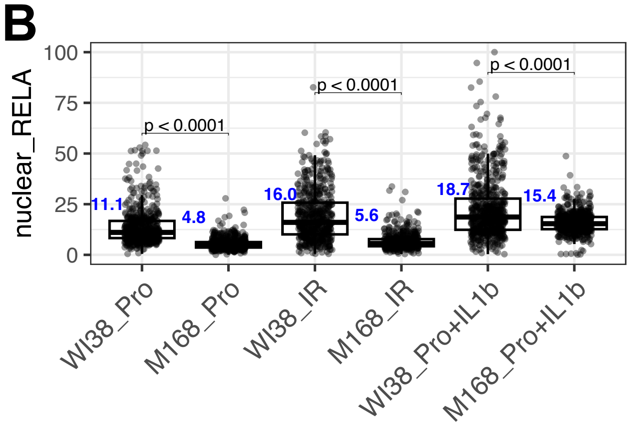


Fig **S6**. Low expression of CXCL8 in M168 cells in IR-induced senescence is correlated with low levels of nuclear RELA, and high expression of CXCL8 is correlated with high levels of nuclear RELA. (A) Immunofluorescence images of RELA and CXCL8 to visualize RELA intracellular localization and CXCL8 levels within WI38 and M168 cells in proliferation (Pro) or after treatment with 10 ng/ml IL1-beta for 2 hours, or 10 days after 40 Gy X-irradation (IR). Nuclear DNA is visualized by staining with DAPI. (**B**) Quantification of RELA nuclear fluorescence. The blue numbers indicate the median of nuclear RELA fluorescence.

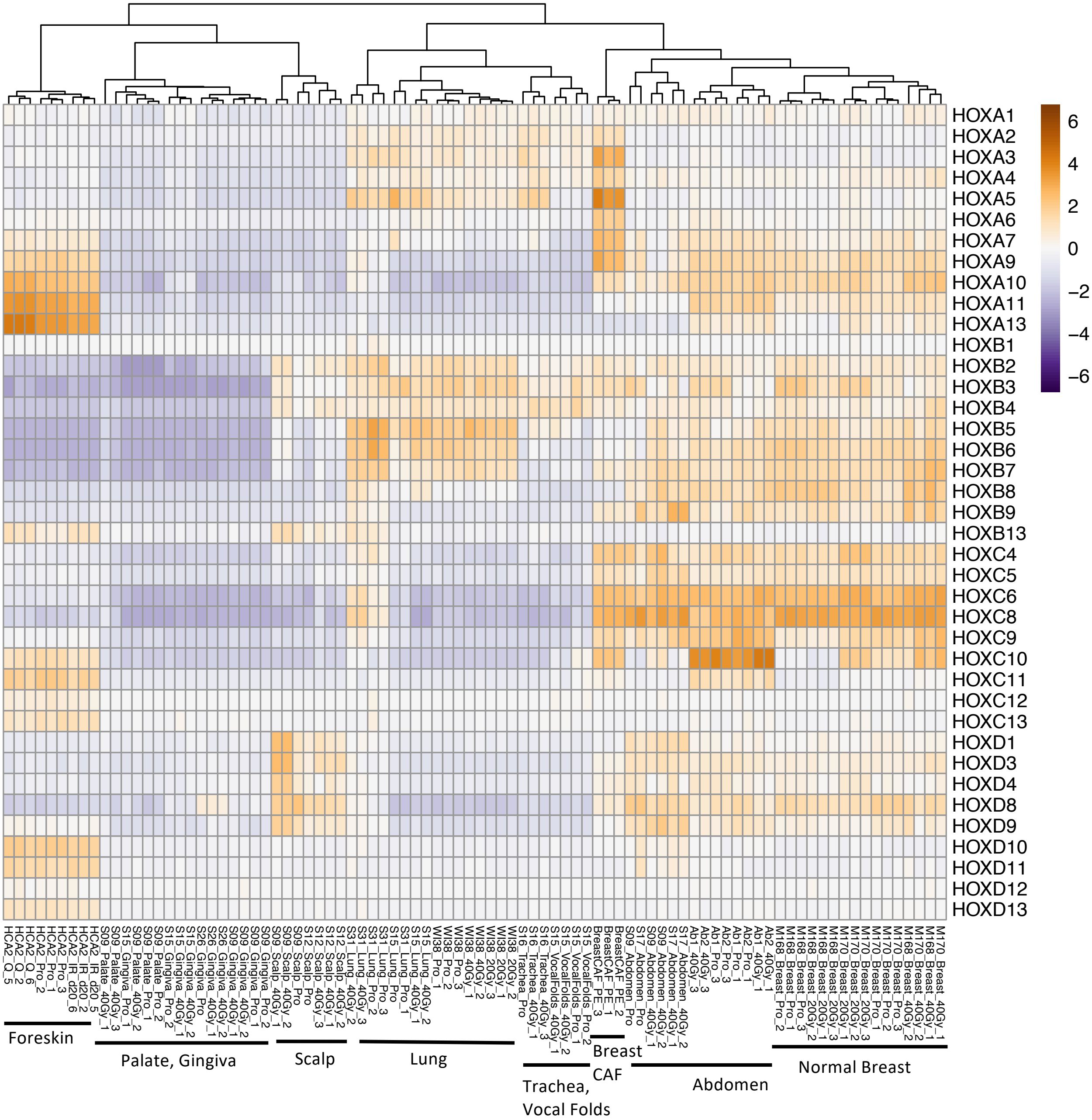


**Fig S7**. HOX gene expression heat map (log2FC) clusters most primary adult fibroblasts by their tissue origin with the exception of Breast CAFs that cluster away from normal breast, and one gingival sample that clusters with palate fibroblasts. HOX gene expression reflects the anatomical position of the tissues. Note that fibroblasts from the gingiva and palate express low levels of all HOX genes, but high levels of some non-HOX homeobox transcription factor genes (Fig. S8). Normal breast and abdominal fibroblasts express high levels of posterior HOX genes. The Breast CAF samples in this study differ from normal breast fibroblasts by the strong expression of some anterior HOXA genes.

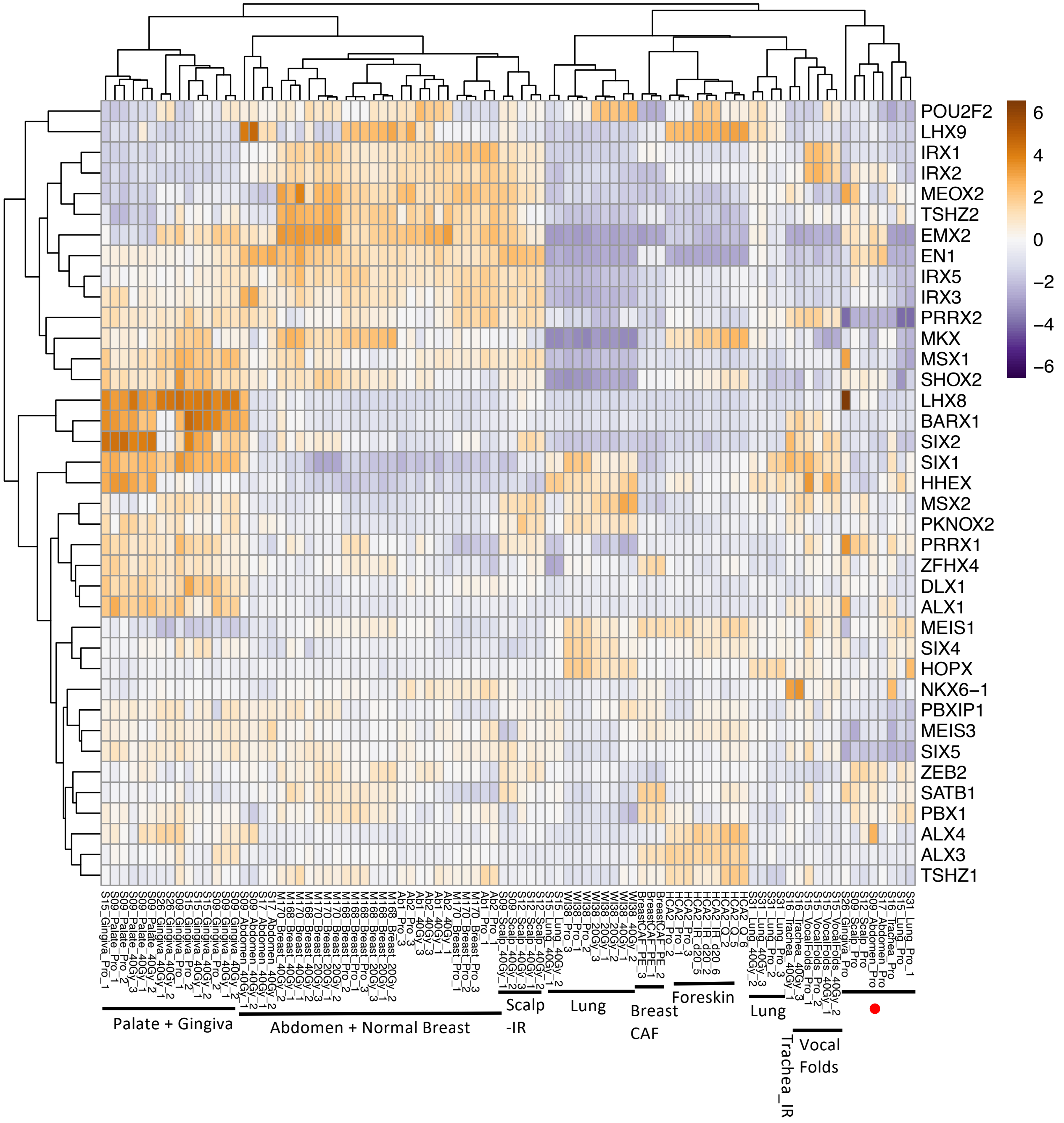


**Fig S8**. Heat map of Non-HOX Homeobox gene expression (log2FC) allows clustering of many fibroblasts by their anatomical position, but less well than the HOX gene heat map. Fibroblasts from the palate and gingiva that expressed all HOX genes at a low level (Fig S7) are seen here to express a characteristic set of non-HOX homeobox transcription factors at a high level. The expression of some Homeobox genes is modified by irradiation leading to a differential clustering of some tissue fibroblasts into proliferation versus IR clusters. Red dot shows a mixed cluster of non-irradiated cells from abdomen, lung, gingiva, and scalp.

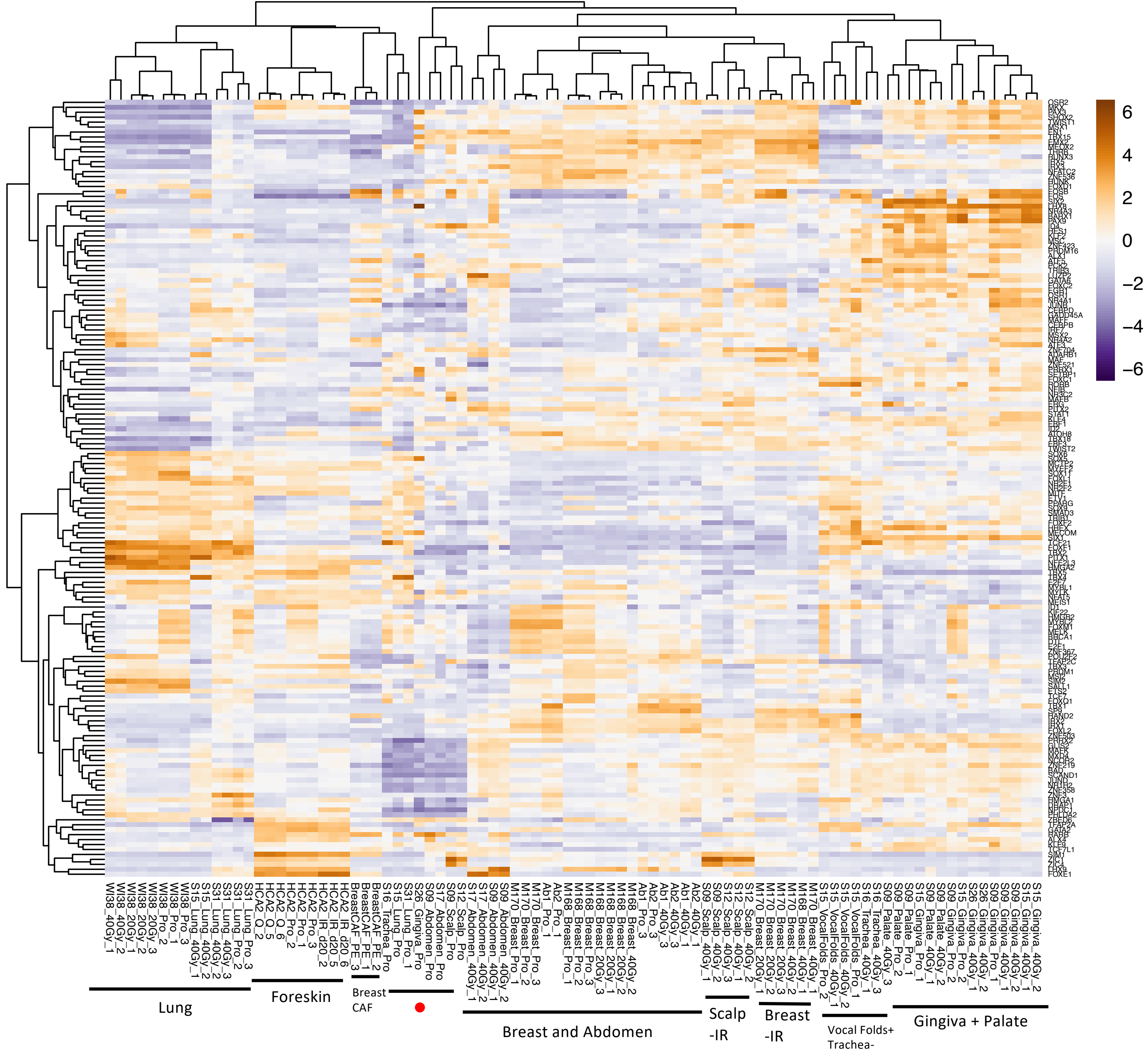


**Fig S9**. Gene expression heat map (log2FC) of the most variable transcription factor genes allows clustering of many fibroblasts by their tissue origin, but less well than the HOX gene heat map. The expression of some transcription factor genes is modified by irradiation leading to a differential clustering of some tissue fibroblasts into proliferation versus IR clusters. The red dot indicates a cluster of proliferating samples from heterogenous tissues.

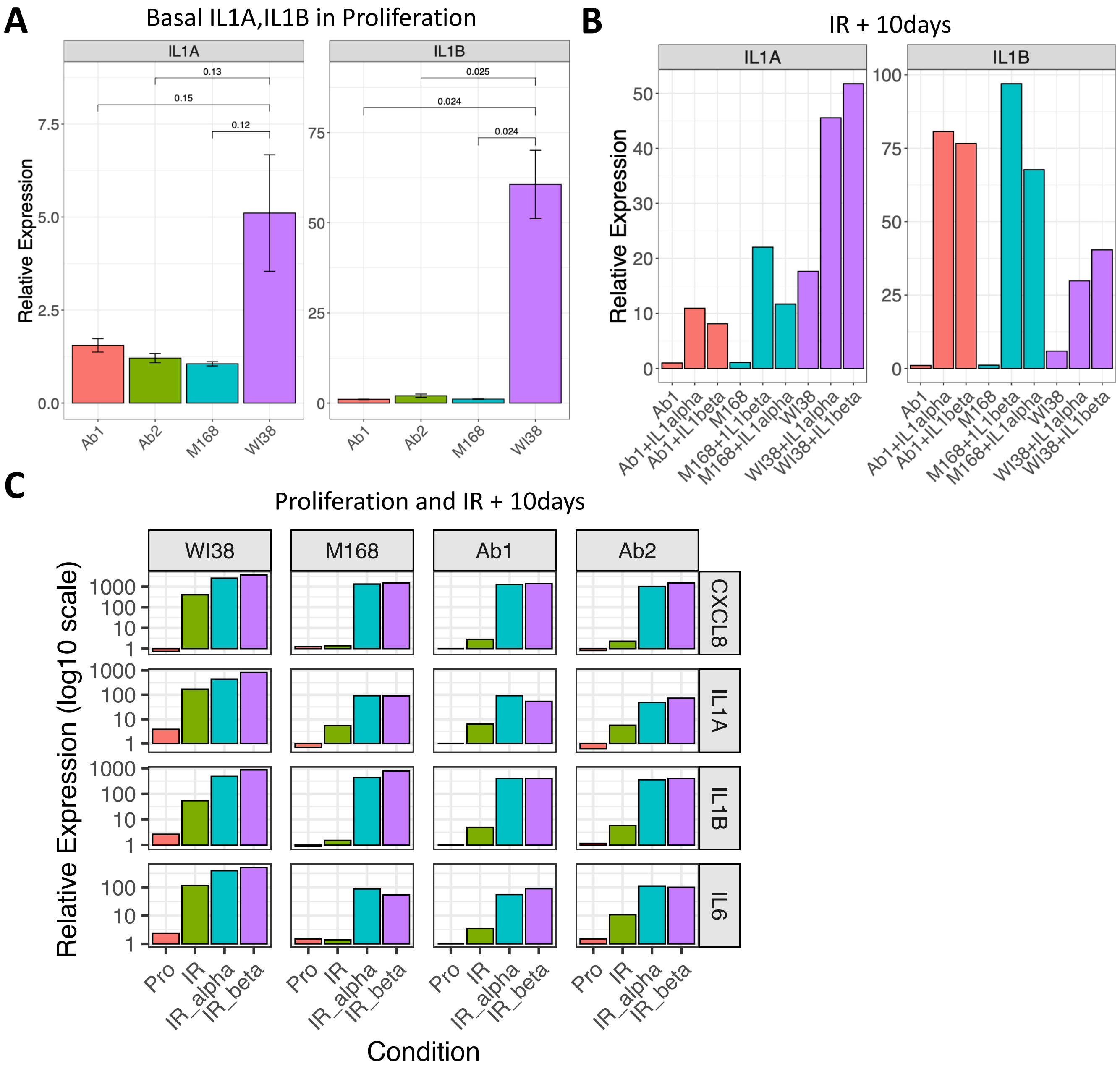


**Fig S10**. RT-qPCR analyses of IL1A, IL1B, IL6, and CXCL8 in WI38 fetal lung, M168 adult mammary, and Ab1 and Ab2 adult abdominal fibroblasts. (**A**) Basal expression of IL1A and IL1B in proliferating Ab1, Ab2, and M168 cells is lower than in WI38 (mean +/- SEM, n=3, Welch's t-test). (**B**) Ab1, M168, and WI38 cells were irradiated with 40 Gy and then incubated 9 days. Cells were then treated with 0 or 20 pg/ml IL1-alpha or IL1-beta for 1 additional day (n=1). (**C**) IR inducibility of CXCL8, IL1A, IL1B, and IL6 is low in Ab1, Ab2, and M168 cells relative to WI38 at 10 days post-irradiation with 40 Gy X-rays. Addition of IL1-alpha or IL1-beta to 20 pg/ml for 24h at 9 days post-irradiation boosted induction of these genes to levels similar to WI38 cells (n=1).

### A) RT-qPCR proliferating cells IL1-alpha dose response

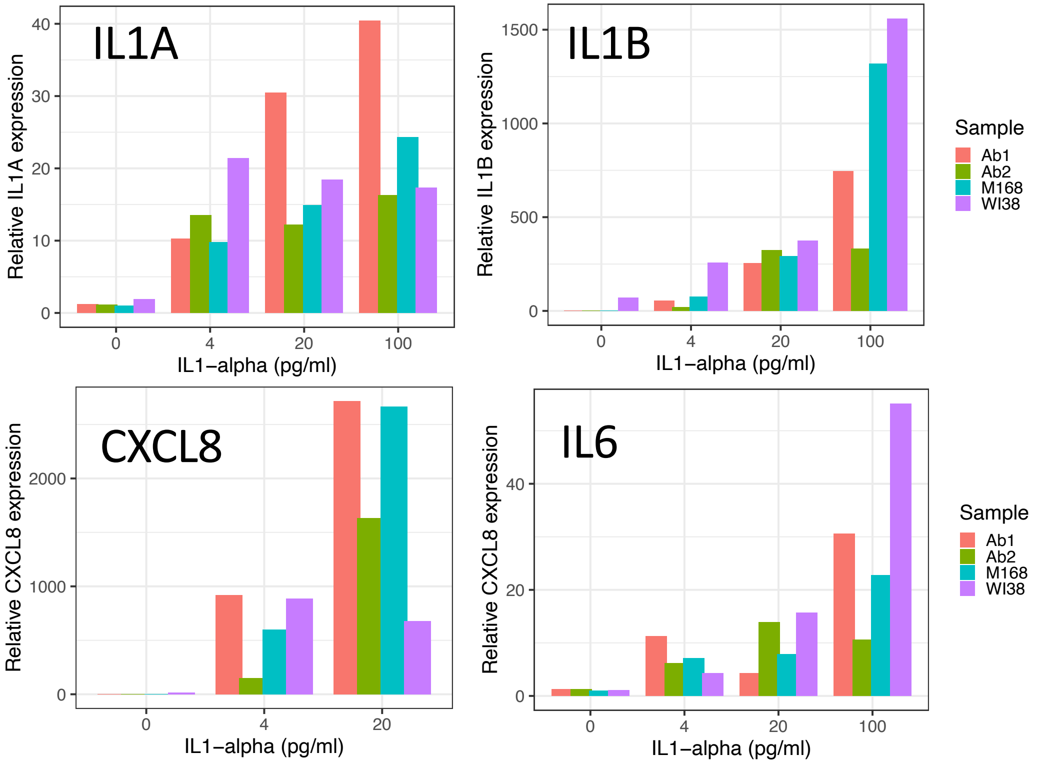


**Fig S11**. Ab1,Ab2,M168, and WI38 cells in proliferation were treated with 0,4,20,or 100 pg/ml of IL1-alpha or IL1-beta for 24 hours and IL1A,IL1B,CXCL8, and IL6 gene expression were quantified by RT-qPCR (n=1).

### B) RT-qPCR proliferating cells IL1-beta dose response

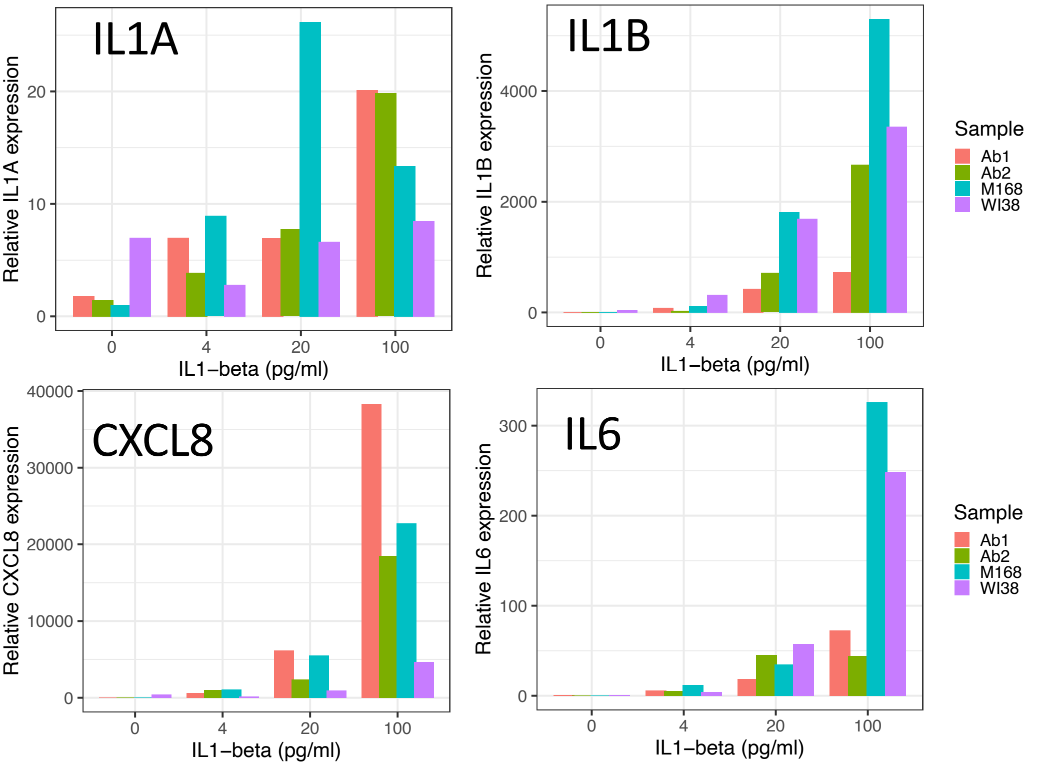

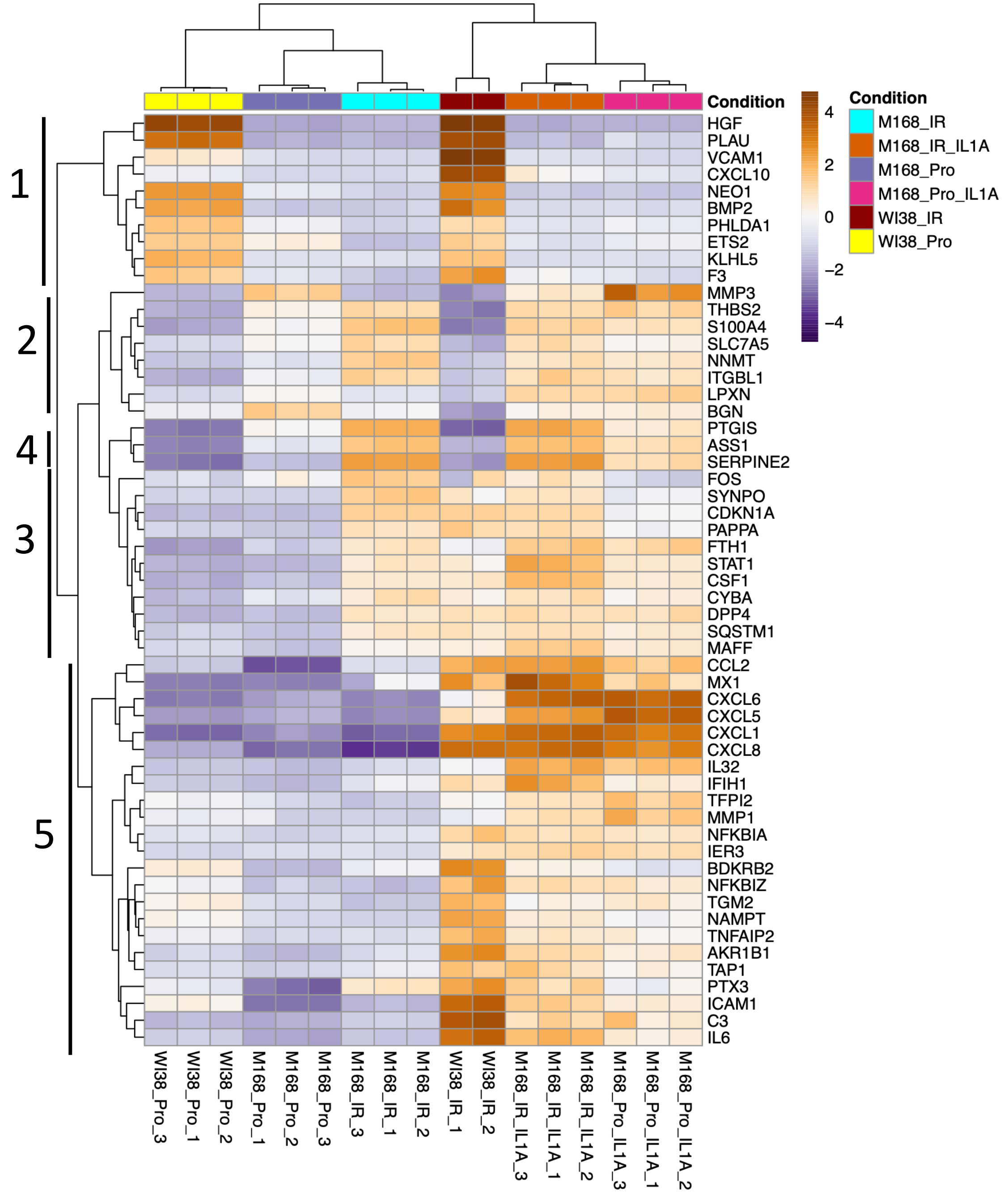


**Fig S12**. Gene expression heat map (log2FC) of 55 RELA-target genes that are highly expressed in either M168 or WI38 cells and that show high expression variance in the experimental conditions comparing both cell lines -/+ IR and -/+ treatment of M168 cells in proliferation or 9 days post-IR with 20 pg/ml IL1-alpha for 24 hours. The IL1A, IL1B, CXCL2, CXCL3, CSF2, and CCL20 genes shown in Fig 5B are not shown in this heat map because only genes with the highest expression levels were retained for this heat map.

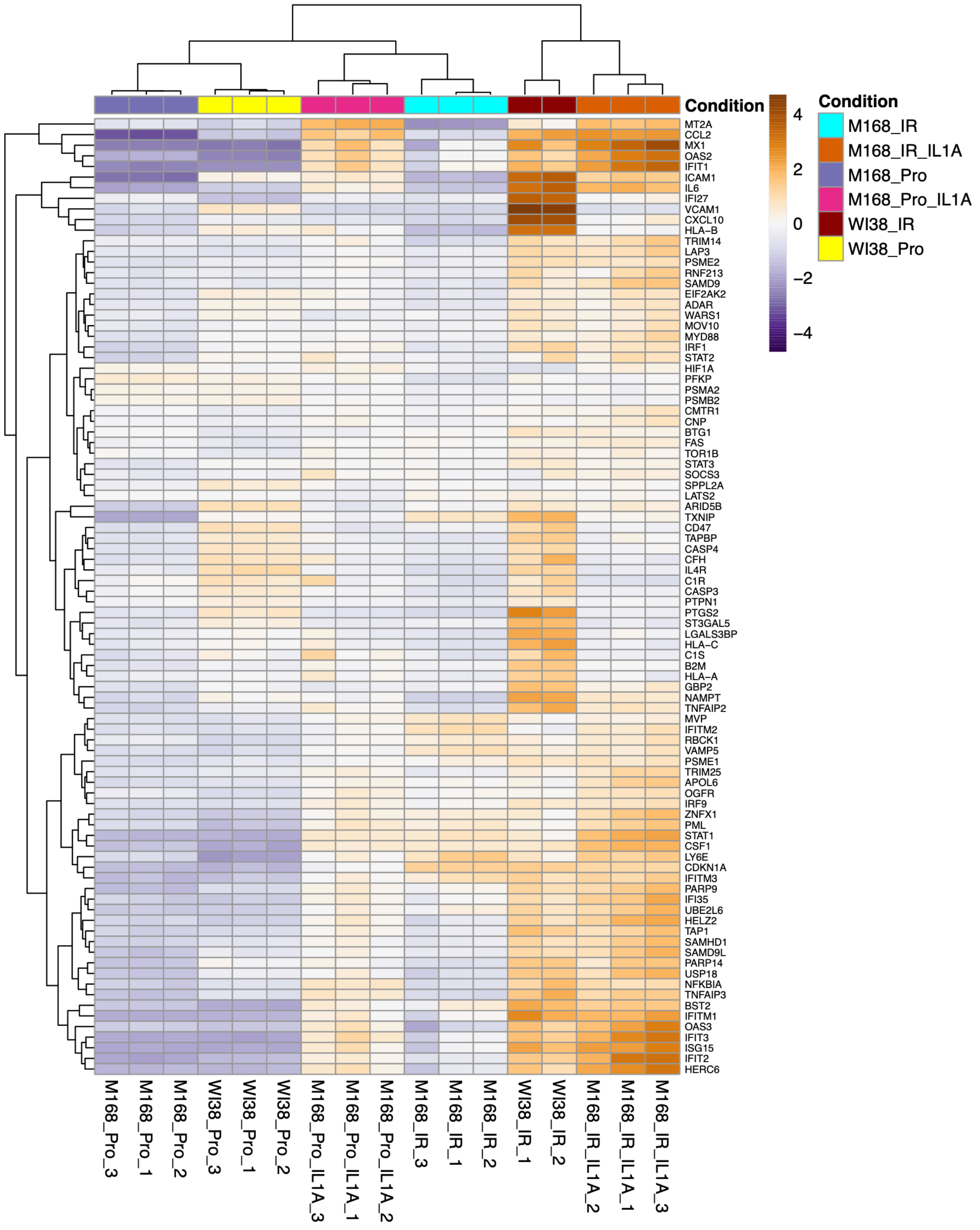


**Fig S13.** Gene expression heat map (log2FC) of 90 ISG (Interferon-Stimulated Genes) that are highly expressed in either M168 or WI38 cells and that show high expression variance in the experimental conditions.

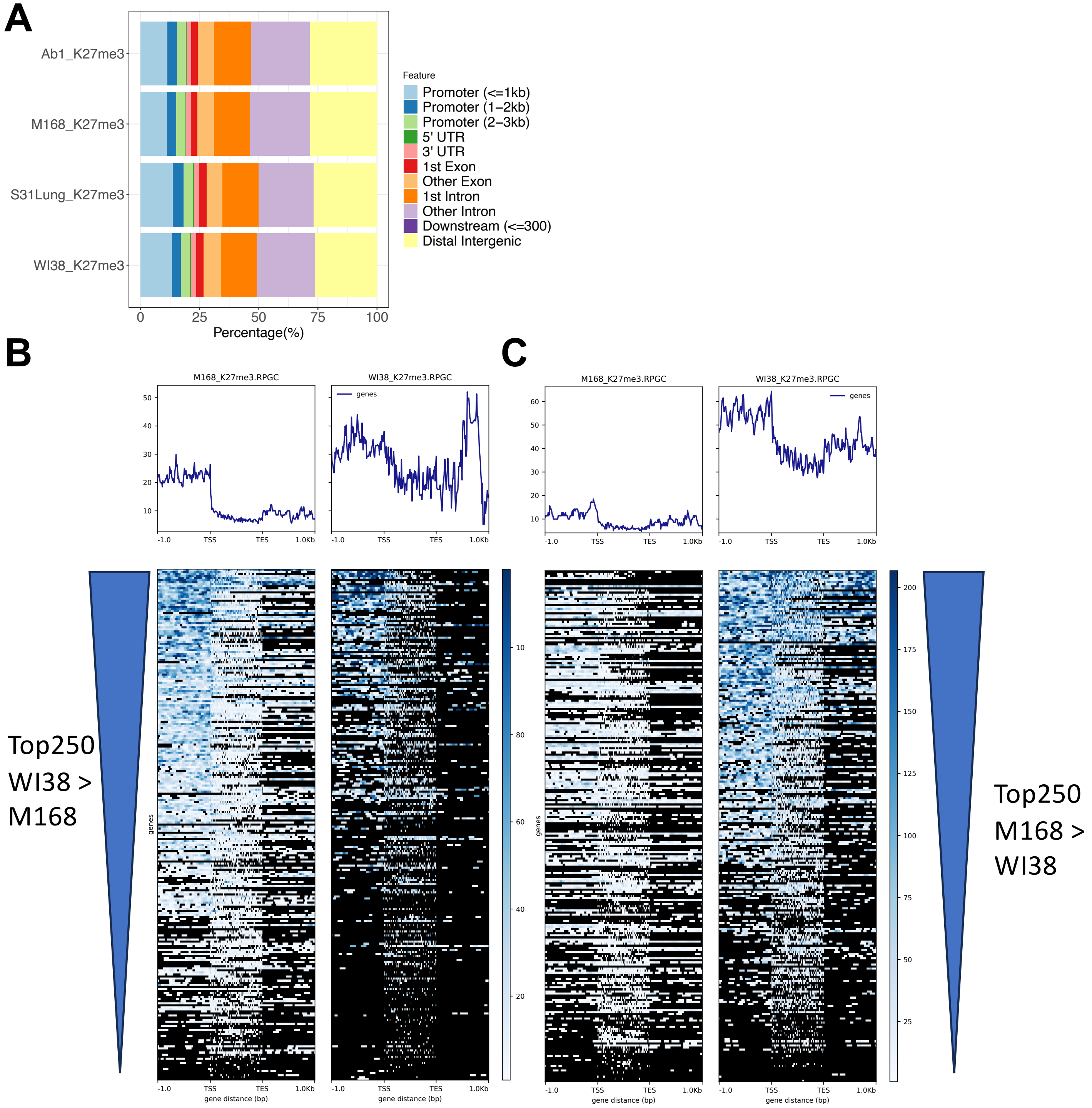


**Fig. S14**. H3-K27me3 analysis. A similar H3-K27me3 genomic distribution is seen for all four primary fibroblasts. (**A**) About 75% of H3-K27me3 is found in the vicinity of genes and about 25% in distal intergenic regions. (**B, C**) H3-K27me3 profile plots (upper panels) and RPGC (reads per genome coverage) heat maps (lower plots) for the top 250 genes that are highly expressed in WI38 versus M168 (B) or in WI38 versus M168 (C). The gene bodies (TSS to TES) were scaled to 1 kb. Overall, high H3-K27me3 density is correlated with cell-type specific repression of gene expression.

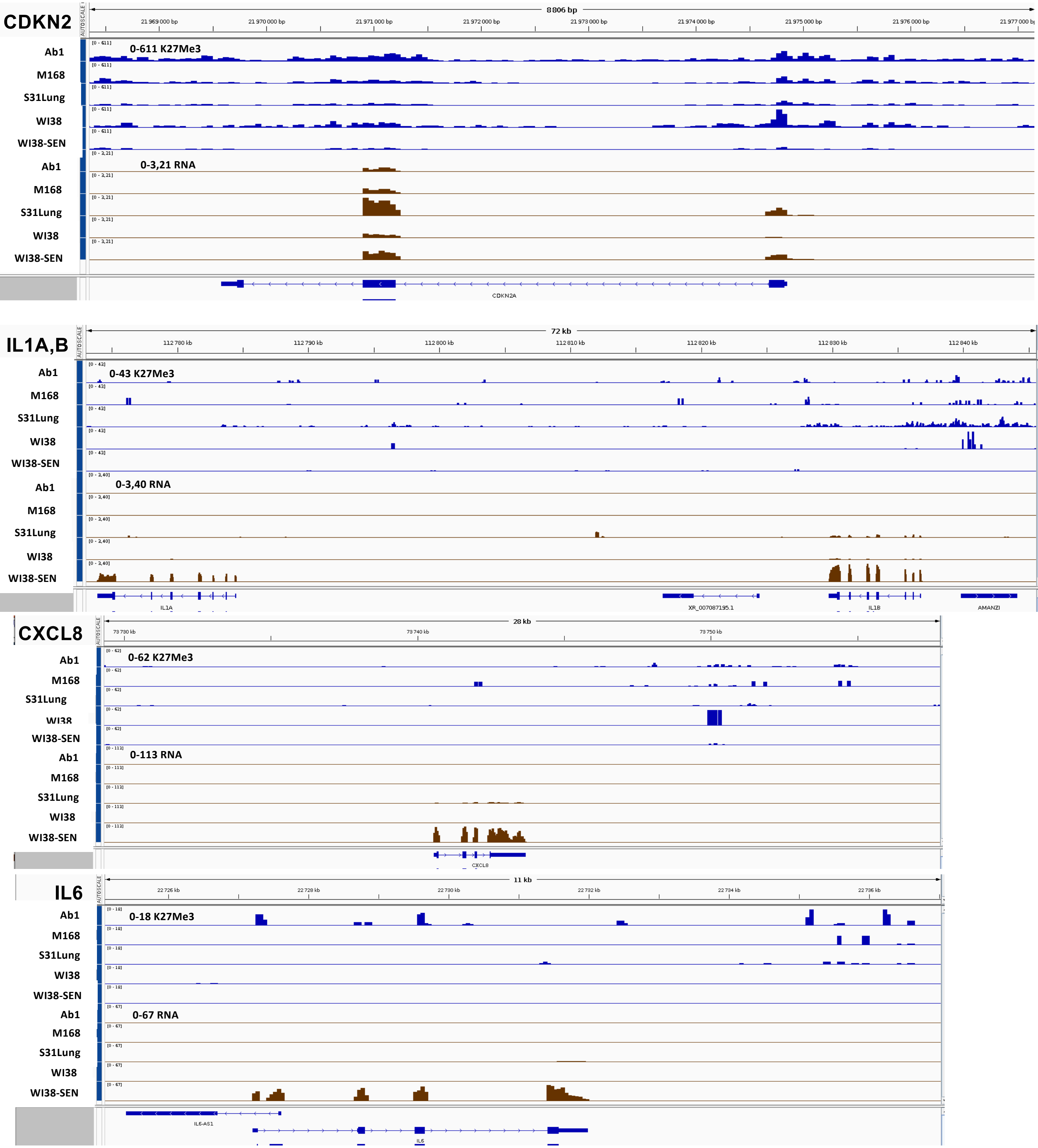


**Fig. S15.** Genome brower images of H3-K27me3 density (RPGC) in blue and mRNA levels (TPM) in brown at the indicated genomic loci.

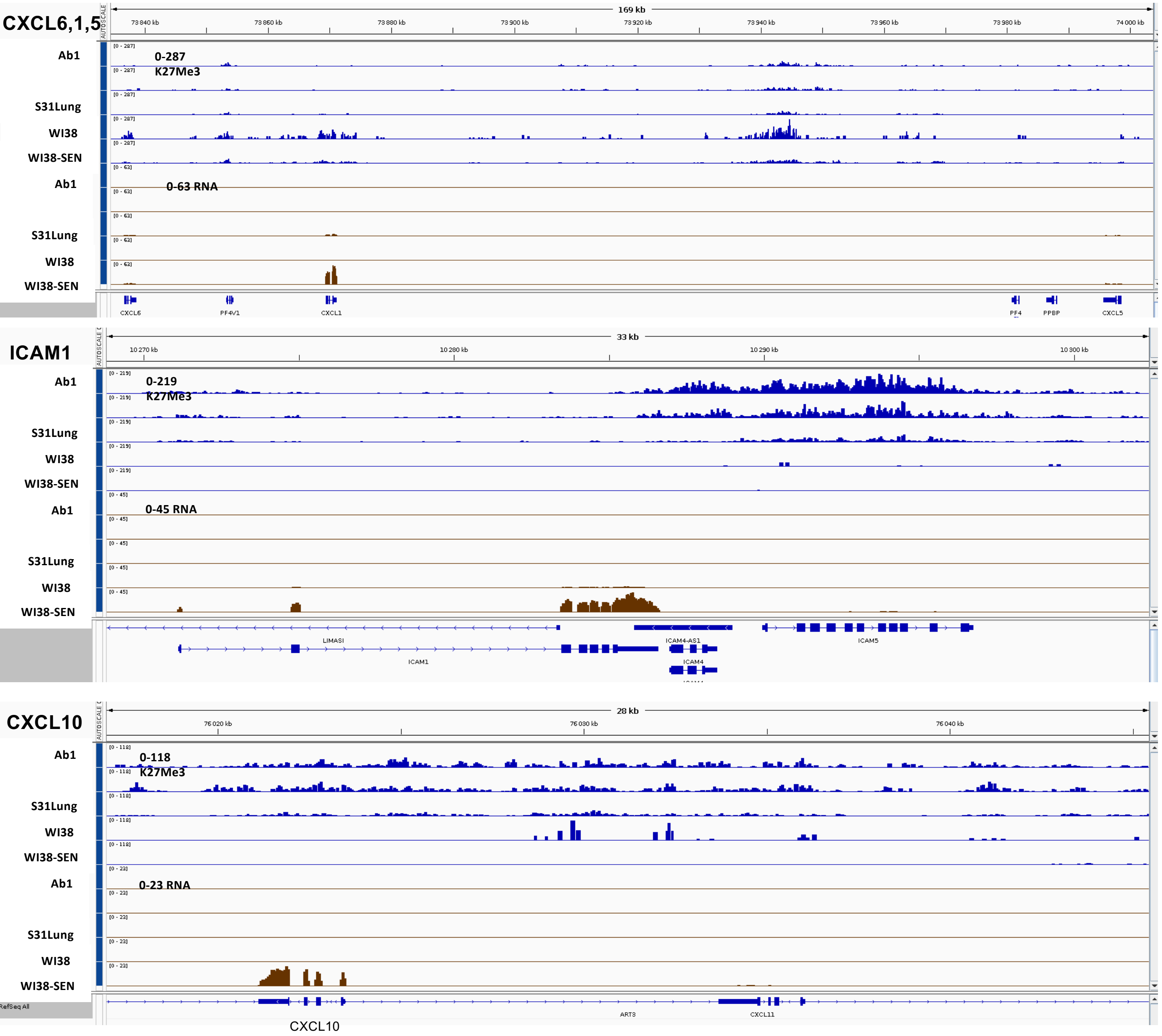


**Fig. S16.** Genome brower images of H3-K27me3 density (RPGC) in blue and mRNA levels (TPM) in brown at the indicated genomic loci.

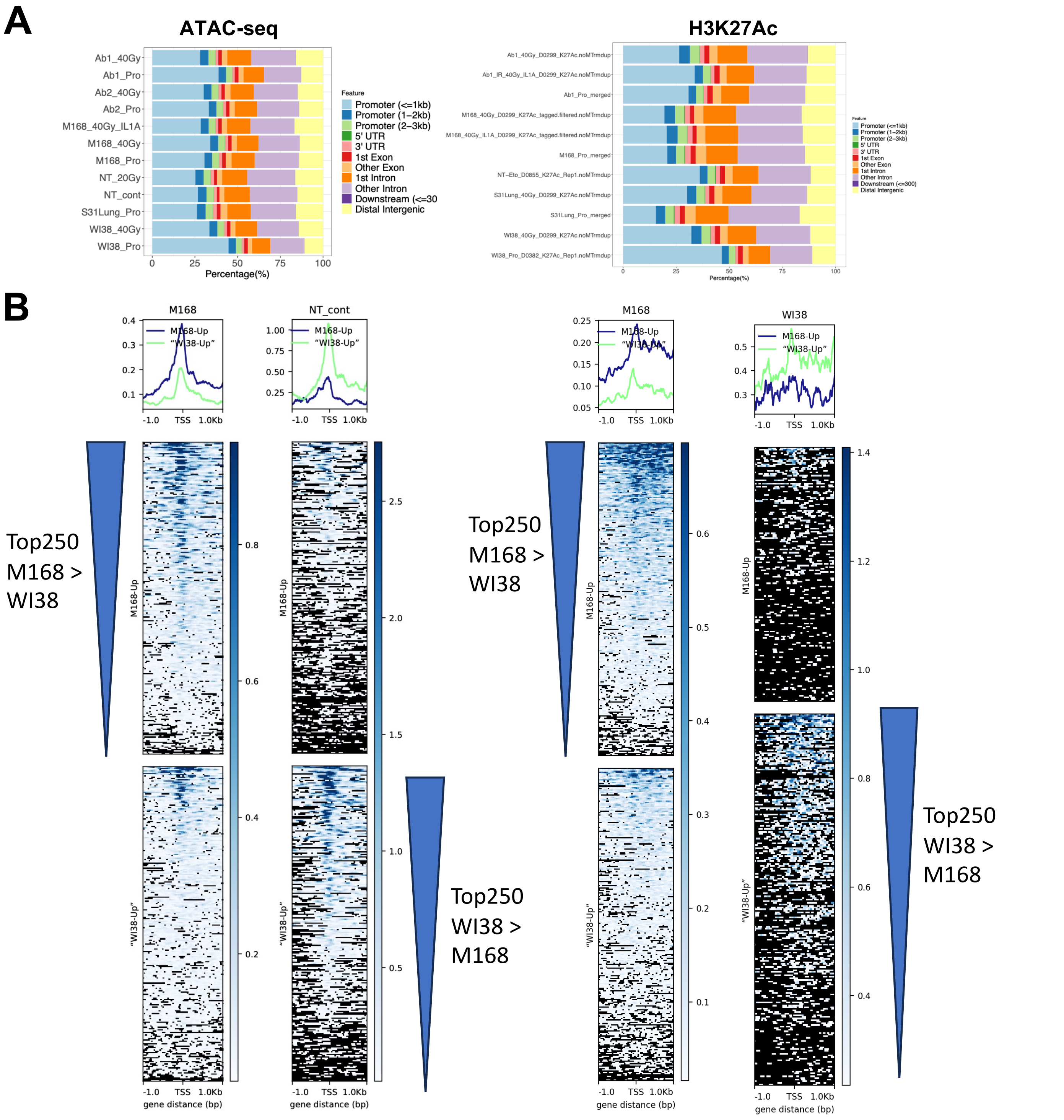


**Fig. S17.** Global ATAC-seq and K3-K27Ac. (**A**) Genomic distributions are similar for ATAC-seq and H3K27Ac. (**B**) profile plots (upper panels) and heat maps (lower plots)

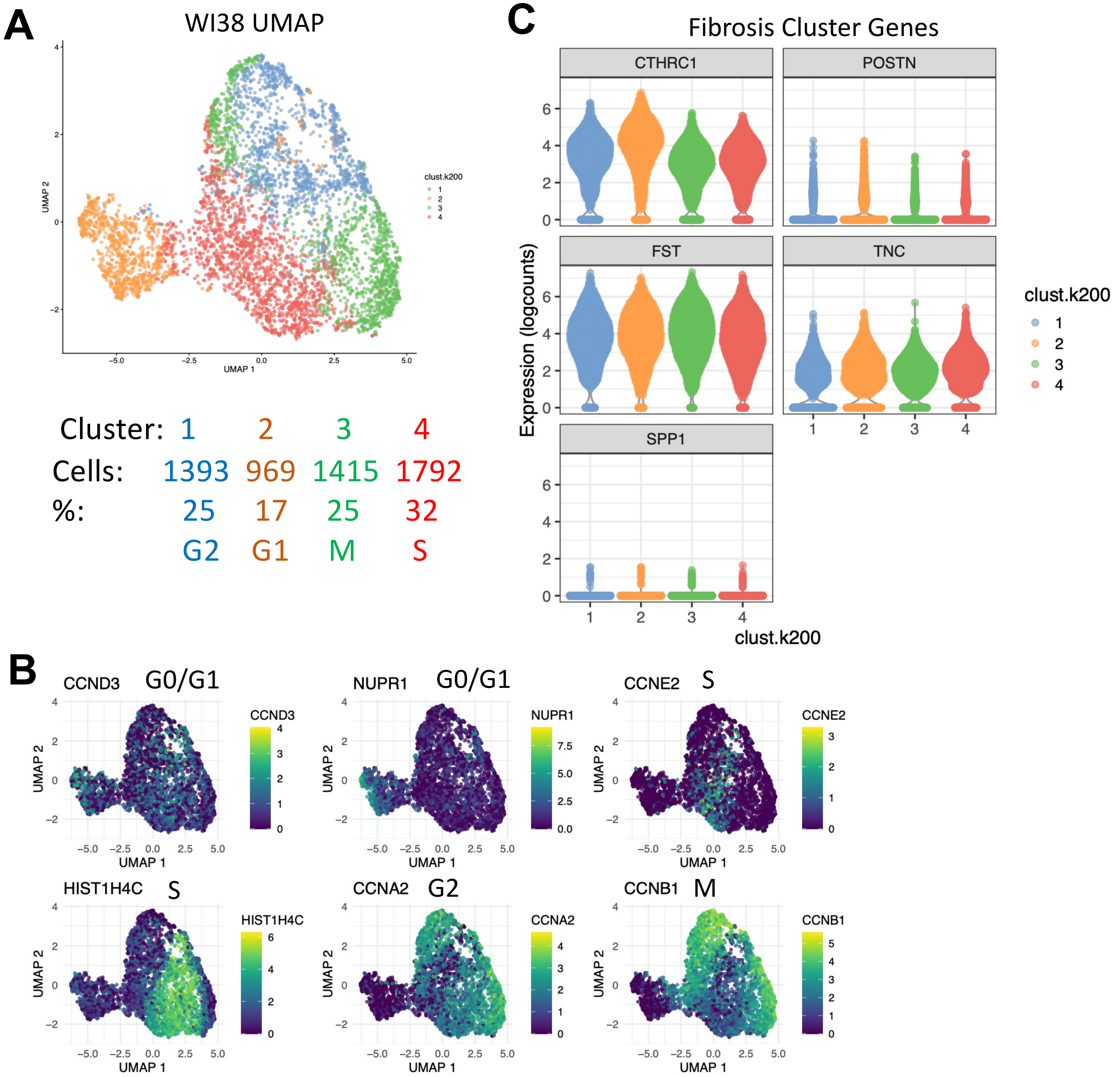


**Fig S18.** Single-cell RNA-seq analysis of proliferating WI38 fibroblasts indicating that these cells manifest a homogeneous transcriptomic profile most similar to fibrotic lung fibroblasts. (**A**) UMAP representation of 4 cell clusters. (**B**) Gene expression heat maps of the indicated cell cycle genes showing that transcriptomic clusters in proliferating WI38 cells correlate with the indicated cell cycle marker genes. It thus appears that the major transcriptomic diffences in proliferating WI38 cells are due to cell cycle phases rather than a mix of different fibroblastic cell types. (**C**) WI38 cells express at moderate to high levels genes that are characteristic of fibrotic lung fibroblasts (CTHRC1, FST, TNC, and POSTN) with the exception of SPP1. Fibrotic fibroblasts are associated with wound healing. WI38 fibroblasts are derived from a fetal lung explant, so their fibrotic profile may be due to preferential migration and amplification of this fibroblastic cell type in the lung explant. Alternatively, the expansion of fibroblasts in vitro in the presence of medium with high levels of serum factors may induce a gene expression program in fibroblasts in vitro that is similar to the fibrotic class of fibroblasts in vivo.

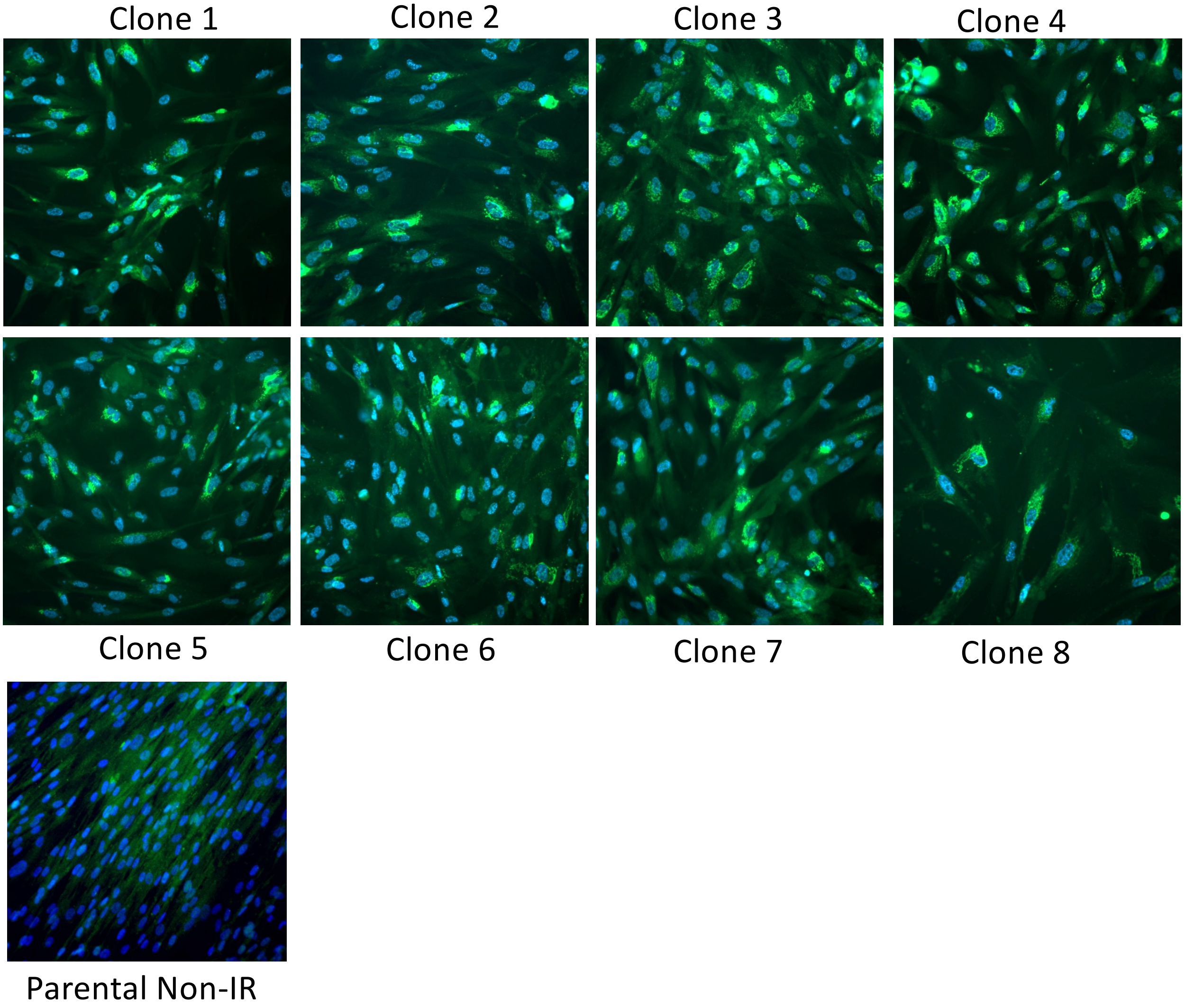


**Fig S19.** WI38hT/DDCas9 clones are competent to induce expression of CXCL8 at 10 days post-irradiation with 40 Gy X-rays. Shown are anti-CXCL8 immuno-fluorescent images (green) with DAPI-stained nuclear DNA (blue) of 8 irradiated clones and the parental non-IR control. All 8 clones showed induction of CXCL8 expression seen within the secretory pathway of positive cells. All clones showed variable percentages of positive cells (40-80%) as for the parental cells after irradiation (Fig 1). We conclude that the parental WI38hT/DDCas9 are not composed of a significant fraction of cells that are refractory to X-ray induced expression of CXCL8.

#### A IL1-Enh2Δ PCR screening

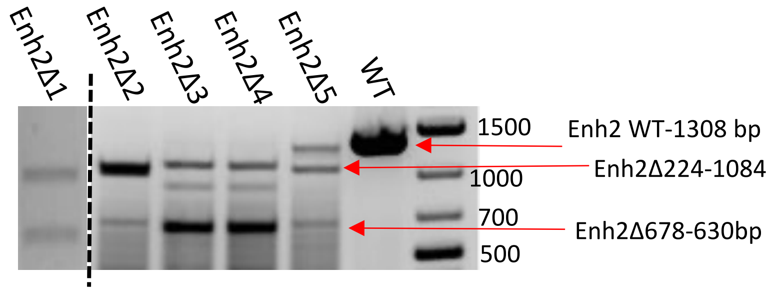


#### B IL1-Enh3Δ PCR screening

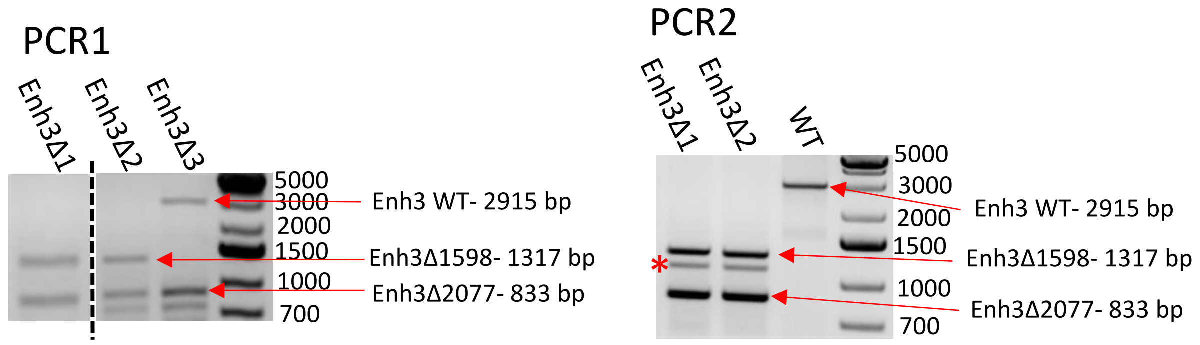


**Fig S20.** IL1-Enh2Δ and Enh3Δ screening by PCR sequencing. (**A**) IL1-Enh2Δ: WI38hTERT/DDCas9 cell were transfected with synthetic guide RNAs Enh2-3 and Enh2-4 (Fig 7A and Table S) to create a 630bp deletion (Enh2Δ630) of one allele. Sanger sequencing indicated that the second allele contained microdeletions at the gRNA cut sites, so guide RNAs Enh2-1 and Enh2-2 targeting an internal region of this second allele were used to delete 224 bp from this second allele. PCR screening showed that the Enh2Δ1 and Enh2Δ2 clones had the expected combination of Enh2Δ630 and Enh2Δ224 alleles. (**B**) IL1-Enh3Δ: WI38hTERT/DDCas9 cell were transfected with synthetic guide RNAs Enh3-1 and Enh3-2 to generate a 2077 bp deletion (Enh3Δ2077) of one allele. Sanger sequencing indicated that the second allele contained microdeletions at the gRNA cut sites, so guide RNAs Enh3-3 and Enh3-4 targeting an internal region of this second allele were used to delete 1598 bp from this second allele. PCR screening showed that the Enh3Δ1 and Enh3Δ2 clones had the expected combination of Enh3Δ2077 and Enh3Δ1598 alleles. Two PCR analyses are shown. The red asterisk in PCR2 indicates a band that was not reproducibly observed (compare with PCR1).

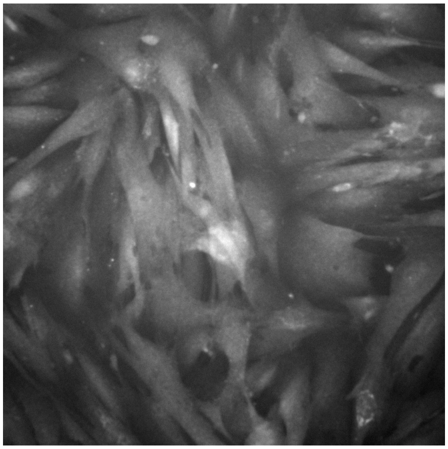

IL1Enh2Δ-IR

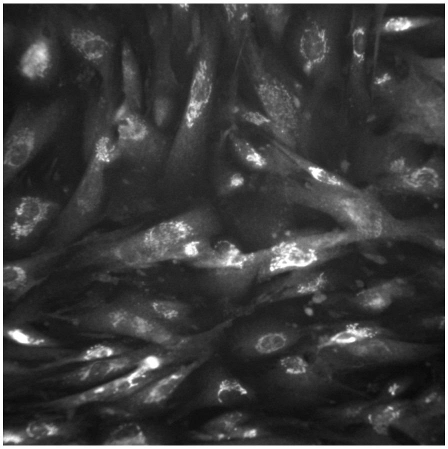

IL1Enh2Δ IR + IL1-beta

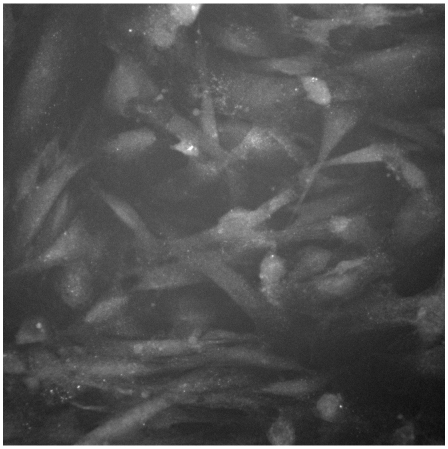

IL1Enh3Δ-IR

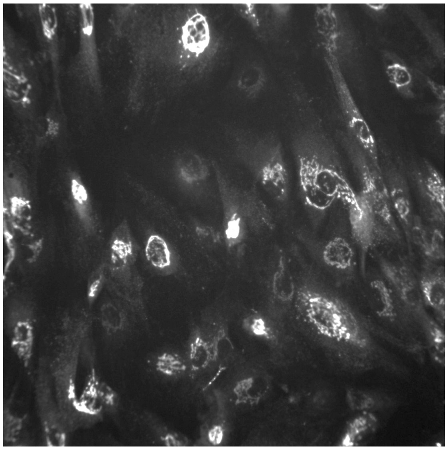

IL1Enh3Δ IR + IL1-beta

**Fig S21.** Absence of CXCL8 expression of CXCL8 in WI38hT cells deleted for IL1-Enh2 or IL1-Enh3 10 days after irradiation with 40 Gy (IR), and strong induction of CXCL8 in the irradiated cells upon addition of IL1-beta to 20 pg/ml and further incubation for 18 hours. The IL1-Enh deletions prevent CXCL8 induction during IR-induced senescence, but they do not prevent induction by a strong stimulation of the RELA pathway by an excess of exogenous IL1-beta.

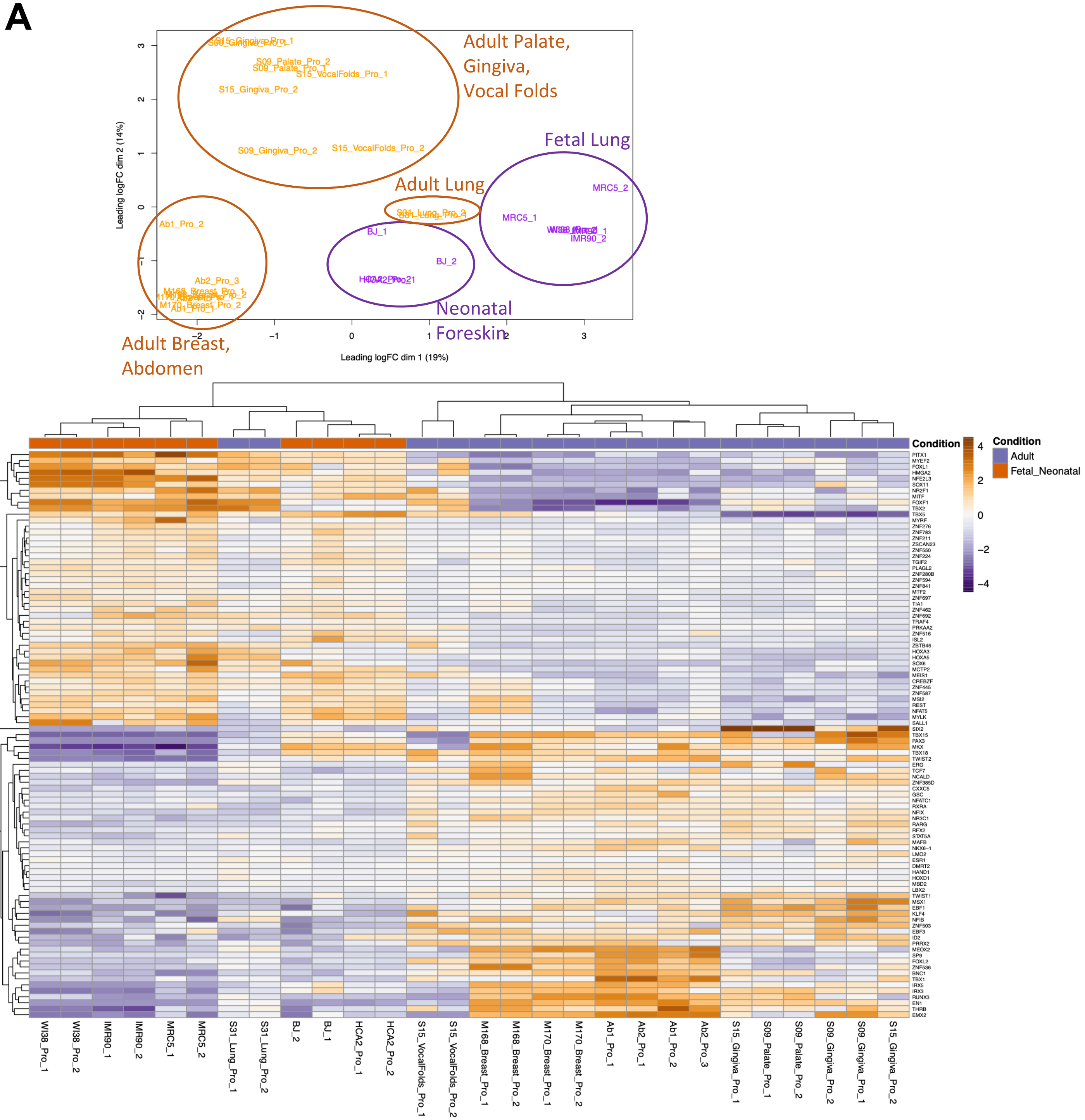


**Fig S22.** Transcription factors that are differentially expressed in fetal/neonatal versus adult fibroblasts. (**A**) Multidimensional scaling showing partial separation of fetal lung/neonatal foreskin versus adult fibroblast transcriptomes. (**B**) Heat map of log2FC expression differences for 46 transcription factor genes showing differential expression between fetal/neonatal versus adult fibroblasts. Note that S31 adult lung fibroblasts share higher similarity to fetal lung/neonatal foreskin fibroblasts for a subset of these transcription factors and thus cluster with the fetal/neonatal fibroblasts. The S31 adult lung fibroblasts also showed intermediate levels of inflammatory gene expression in IR-SEN (Fig. 4A).

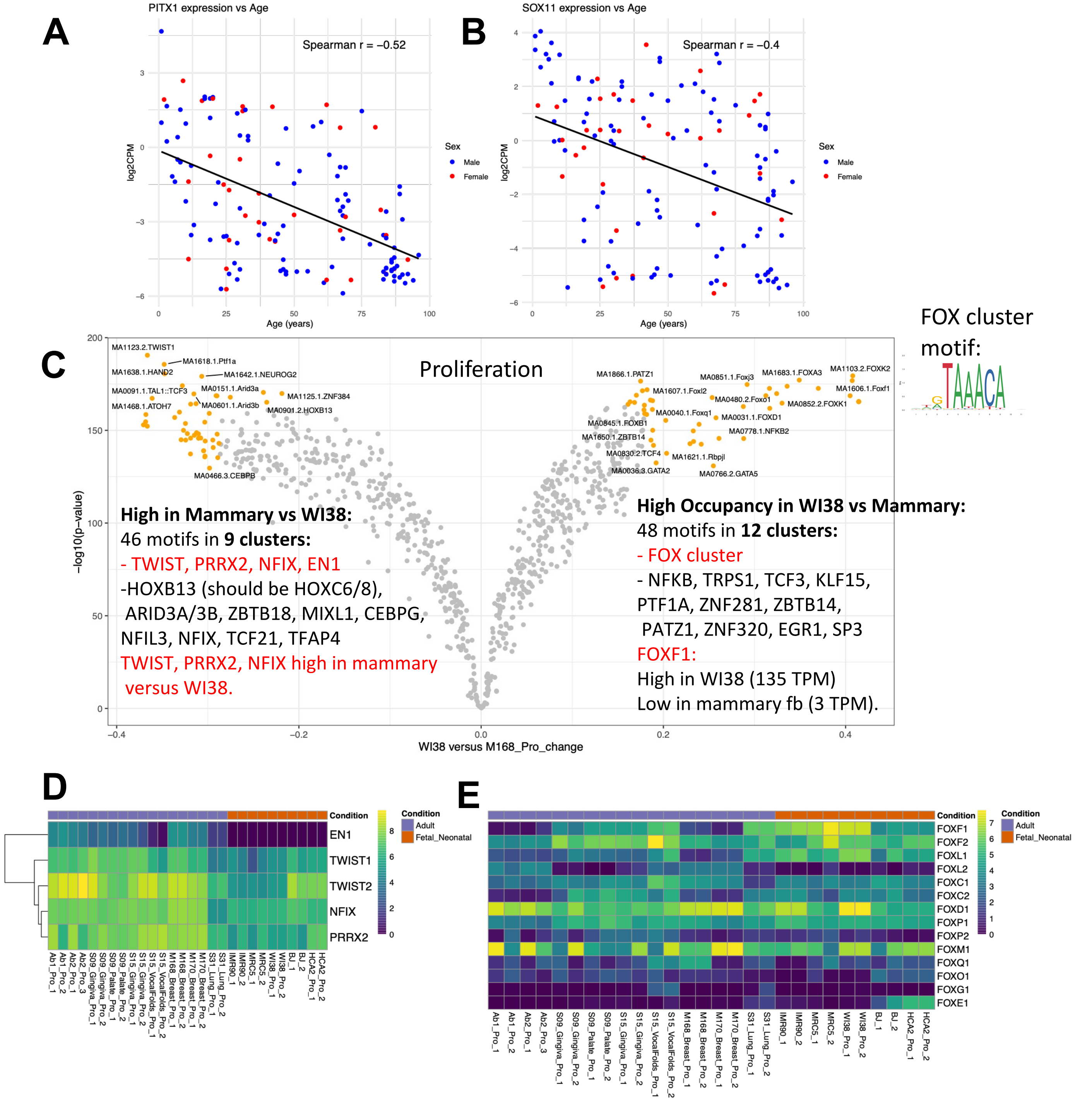


**Fig S23.** Transcription factors differentially expressed in fetal/neonatal versus adult fibroblasts. PITX1 (**A**) and SOX11 (**B**) expression are inversely correlated with the donor age of dermal fibroblasts. (**C**) Differential transcription factor occupancy for proliferating fetal lung WI38 versus adult mammary M168 cells from ATAC-seq data as predicted by the TOBIAS software package. (**D,E**) Log2FC gene expression heat maps for transcription factor genes expressed higher in adult (**D**) or higher in fetal/neonatal fibroblasts (**E**).

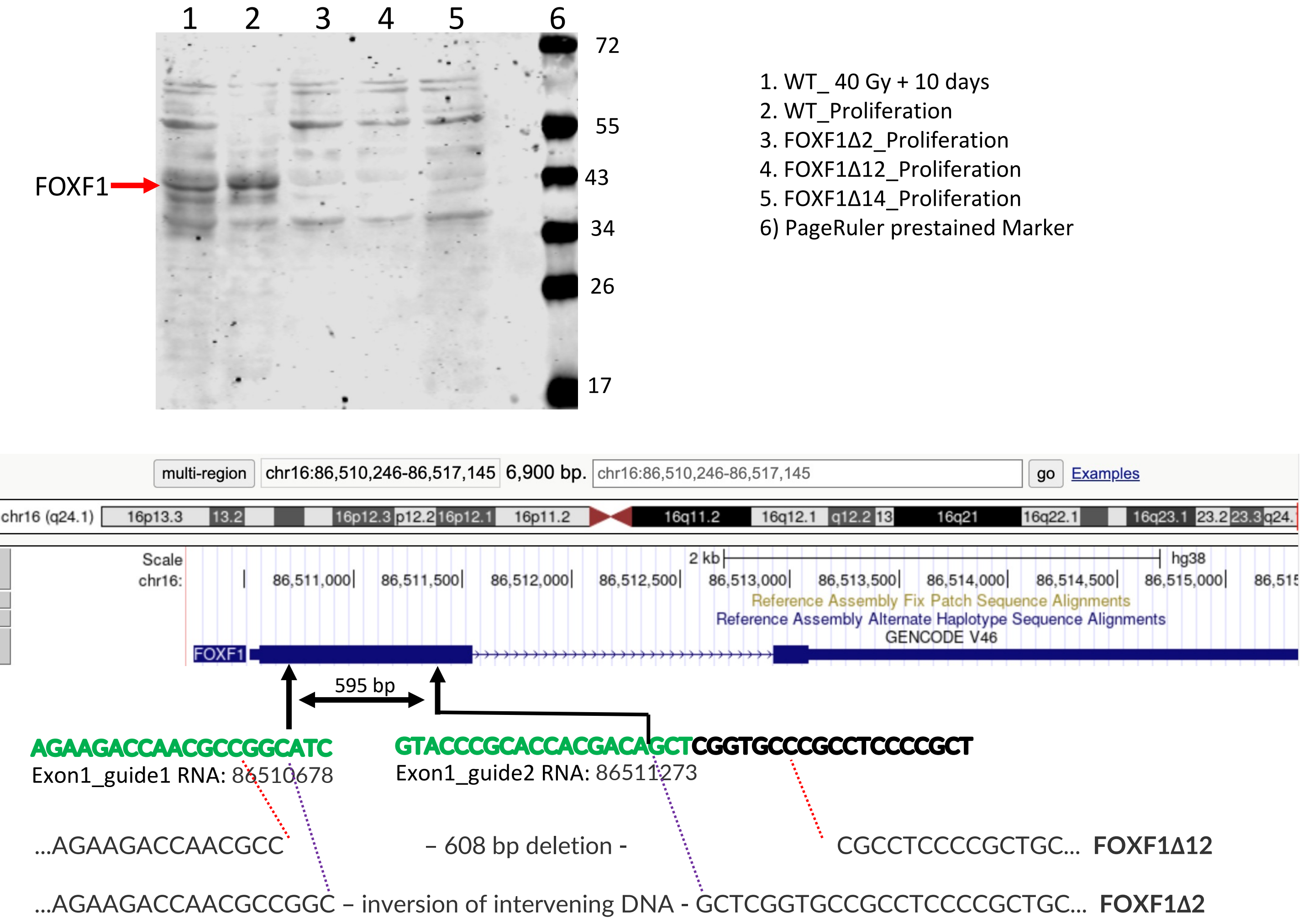


**Fig S24.** Analysis of FOXF1Δ2 and FOXF1Δ12 knock-out clones. **(A)** FOXF1 Western blots showing the apparent absence of FOXF1 expression in the FOXF1Δ2, FOXF1Δ12, and FOXF1Δ14 clones relative parental WI38hTERT cells. FOXF1 levels were similar in irradiated and proliferating WT cells. (B) KO deletion breakpoints determined by the analysis of short-read DNA sequences spanning the Guide1 and Guide2 (green sequences) RNA-directed cut sites. The FOXF1Δ12 reads showed a 608bp deletion encompassing the Guide1 and Guide2 cut sites that would delete 203 aa including the DNA binding domain of the protein and lead to a truncated protein. The FOXF1Δ2 DNA sequences showed breakpoints consistent with cutting at the predicted Guide1 and Guide2 cut sites with an inversion and religation of the intervening DNA sequence leading to the expression of a frameshifted protein after 42 aa with a premature stop codon.
